# RNA-binding proteins provide specificity to the PAN2–PAN3 mRNA deadenylation complex

**DOI:** 10.1101/2025.09.27.678968

**Authors:** Terence T.L. Tang, Francesco Zangari, Conny W.H. Yu, James A.W. Stowell, Alžběta Roeselová, Stefan M.V. Freund, Anne-Claude Gingras, Lori A. Passmore

## Abstract

Cytoplasmic shortening of mRNA poly(A) tails represses eukaryotic gene expression by inhibiting efficient translation and committing an mRNA to decay. The CCR4–NOT deadenylase machinery interacts with sequence-specific RNA-binding proteins (RBPs), termed RNA adaptors, to target specific transcripts for deadenylation. In contrast, the PAN2–PAN3 deadenylation complex is thought to be predominantly recruited to mRNAs via interaction with the poly(A) binding protein, raising the question of whether it acts in a transcript-specific manner. Here, using biochemical reconstitution, we show that PAN2–PAN3 can also be recruited to specific RNAs via RNA adaptors, including MEX3, YTHDF and ZFP36 proteins. In cells, we find that a diverse range of RNA adaptors interact with both major deadenylation complexes. Thus, our data suggest that, in addition to CCR4-NOT, PAN2-PAN3 also contributes to the specificity of mRNA degradation and the robustness of post-transcriptional regulation of gene expression.

## Introduction

The abundance of cytoplasmic transcripts in eukaryotes is controlled by mRNA synthesis and decay rates to regulate gene expression. The half-life of an mRNA can range from minutes to hours (Friedel et al. 2009; Geisberg et al. 2014; Wang et al. 2002). Canonical mRNA decay is initiated by shortening of the mRNA 3′ polyadenosine (poly(A)) tail, or deadenylation (Parker and Song 2004), which is mainly carried out by two highly conserved multiprotein complexes: PAN2–PAN3 and CCR4–NOT (Passmore and Coller 2022). Deadenylation represses gene expression by releasing the cytoplasmic poly(A)-binding protein PABPC1 to reduce translation efficiency, and by promoting decapping and exonucleolytic degradation (Coller and Parker 2004). The importance of deadenylation in gene expression is exemplified by its involvement in many processes, including eukaryotic development (Fox and Wickens 1990), viral RNA stability (Garneau et al. 2008), and the immune system (Hao and Baltimore 2009).

The half-lives of individual eukaryotic transcripts are modulated by specific targeting to the deadenylation machinery. This specificity has largely been attributed to *cis* RNA elements found in 3′ untranslated regions (UTRs), many of which are binding sites for RNA-binding proteins (RBPs) (Cheng et al. 2017; Van Nostrand et al. 2020b). Significant progress has been made in mapping RBP-mRNA interactions (Lin and Miles 2019; Schueler et al. 2014; Silverman et al. 2014; Van Nostrand et al. 2020a), but the exact consequences of these interactions are less well understood. In some cases, RBPs promote deadenylation by acting as RNA adaptors, binding simultaneously to specific RNA motifs and to CCR4–NOT (Passmore and Coller 2022). However, for many RBPs, a mechanistic link to deadenylation specificity and mRNA half-life remains unclear.

RNA-binding proteins are enriched in intrinsically disordered regions (IDRs) - polypeptide segments that lack defined secondary and tertiary structures (Varadi et al. 2015; Wang et al. 2016; Zhao et al. 2021). IDRs can be involved in RNA recognition (Basu and Bahadur 2016) or in interactions with other proteins through diverse mechanisms. For example, sequence-specific RBPs such as Pumilio and TTP, or miRNA-associated proteins such as GW182 (TNRC6 in metazoans), interact with the CCR4–NOT deadenylase complex (Arvola et al. 2020; Chen et al. 2009; Eulalio et al. 2009; Webster et al. 2019). Interactions are often mediated through a multivalent mechanism involving short-linear motifs (SLiMs) and hydrophobic or aromatic side chains within the IDRs of RNA-binding proteins (Bhandari et al. 2014; Raisch et al. 2016; Sgromo et al. 2017; Stowell et al. 2024; Pekovic et al. 2025). Extended regions of IDRs can interact with multiple sites on CCR4–NOT. Such interactions effectively tether CCR4-NOT to the cognate RNAs to promote targeted deadenylation and mRNA decay.

Similarly, PAN2–PAN3 interacts with GW182 (Christie et al. 2013) to promote deadenylation of transcripts in a miRNA-targeted manner (Braun et al. 2011; Chekulaeva et al. 2011; Fabian et al. 2011). PAN2-PAN3 also co-immunoprecipitates with YTHDF3 to regulate development (Liu et al. 2020), suggesting that it may promote deadenylation of YTHDF3-bound transcripts. However, unlike CCR4–NOT, PAN2–PAN3 is not known to play a major role in targeted deadenylation, and it is not known to directly interact with a large number of RBPs.

PAN2–PAN3 interacts with the poly(A) binding protein PABPC1 (Mangus et al. 2003), recognizing the interfaces between sequential PABPC1 molecules (Schäfer et al. 2019). Depletion of PAN2–PAN3 in cells stabilizes longer poly(A) tails at steady state, whereas depletion of CCR4–NOT stabilizes shorter poly(A) tails (Boeck et al. 1996; Tucker et al. 2001; Webster et al. 2019; Yi et al. 2018; Yamashita et al. 2005). This led to the ‘biphasic model’ of deadenylation where PAN2–PAN3 functions in the shortening of the distal region of long poly(A) tails whereas CCR4–NOT commits a transcript to degradation by removing the proximal part of poly(A) tails in a targeted manner (Wahle and Winkler 2013; Passmore and Coller 2022). The pleotropic effects of interfering with mRNA stability in cells complicates study of this process. Nevertheless, *in vivo* studies have demonstrated functional redundancy between PAN2–PAN3 and CCR4–NOT, suggesting that PAN2–PAN3 may compensate for CCR4–NOT in RBP-mediated targeted deadenylation (Bönisch et al. 2007; Tucker et al. 2001; Yamashita et al. 2005). Thus, we aimed to use a fully defined, reconstituted system to examine the role of PAN2–PAN3.

The prevailing model in the field is that PAN2–PAN3 predominantly acts non-discriminately on long poly(A) tails. We hypothesized that PAN2–PAN3 is highly regulated to provide specificity to mRNA deadenylation. Here, we set out to determine whether PAN2– PAN3 carries out transcript-targeted deadenylation. We discover that functionally diverse RBPs can interact with the human PAN2–PAN3 complex, acting as RNA adaptors that promote targeted deadenylation. Therefore, our results reveal a more pervasive role for specificity in mRNA deadenylation. Taken together, our work provides a potential explanation for the functional redundancy of PAN2–PAN3 and CCR4–NOT that would ensure robust and high-fidelity post-transcriptional regulation of gene expression.

## Results

### Identification of RBP interactors of PAN2–PAN3

Proximal interactors of CCR4–NOT had been previously identified in a proximity-based proteomics (BioID) study performed in HEK293 cells (Youn et al. 2018). CCR4–NOT was shown to cluster with the miRNA-associated machinery, including GW182 (TNRC6A-C), Argonaute (AGO1-3), and GIGYF (GIGYF1-2) proteins. Additionally, CCR4–NOT subunits cluster with RNA adaptors, such as Pumilio (PUM1-2), YTHDF (YTHDF1-3), and Roquin (RC3H1-2) proteins. These interactions are consistent with the miRNA machinery and sequence-specific RBPs contributing to the specificity of deadenylation by CCR4-NOT.

PAN2 and PAN3 had not been tagged with biotin ligase by Youn et al., but they were detected as interactors of other tagged proteins. We further examined the dataset and found that 20 bait proteins are proximal to both PAN2 and PAN3 (**Figure 1a and S1a, Table S1**). These include known interactors of PAN2–PAN3, such as PABPC1 (Uchida et al. 2004) and the TNRC6 proteins (Braun et al. 2011; Huntzinger et al. 2013; Kuzuoglu-Öztürk et al. 2012), but also other RNA-binding proteins that were not known to interact with PAN2– PAN3 (**Figure 1a**). We sorted the hits by the number of identified PAN2 or PAN3 peptides, and found that the MEX3B, YTHDF2, YTHDF3 and ZFP36 RBPs recover the highest relative amounts of both proteins.

**Figure 1.**
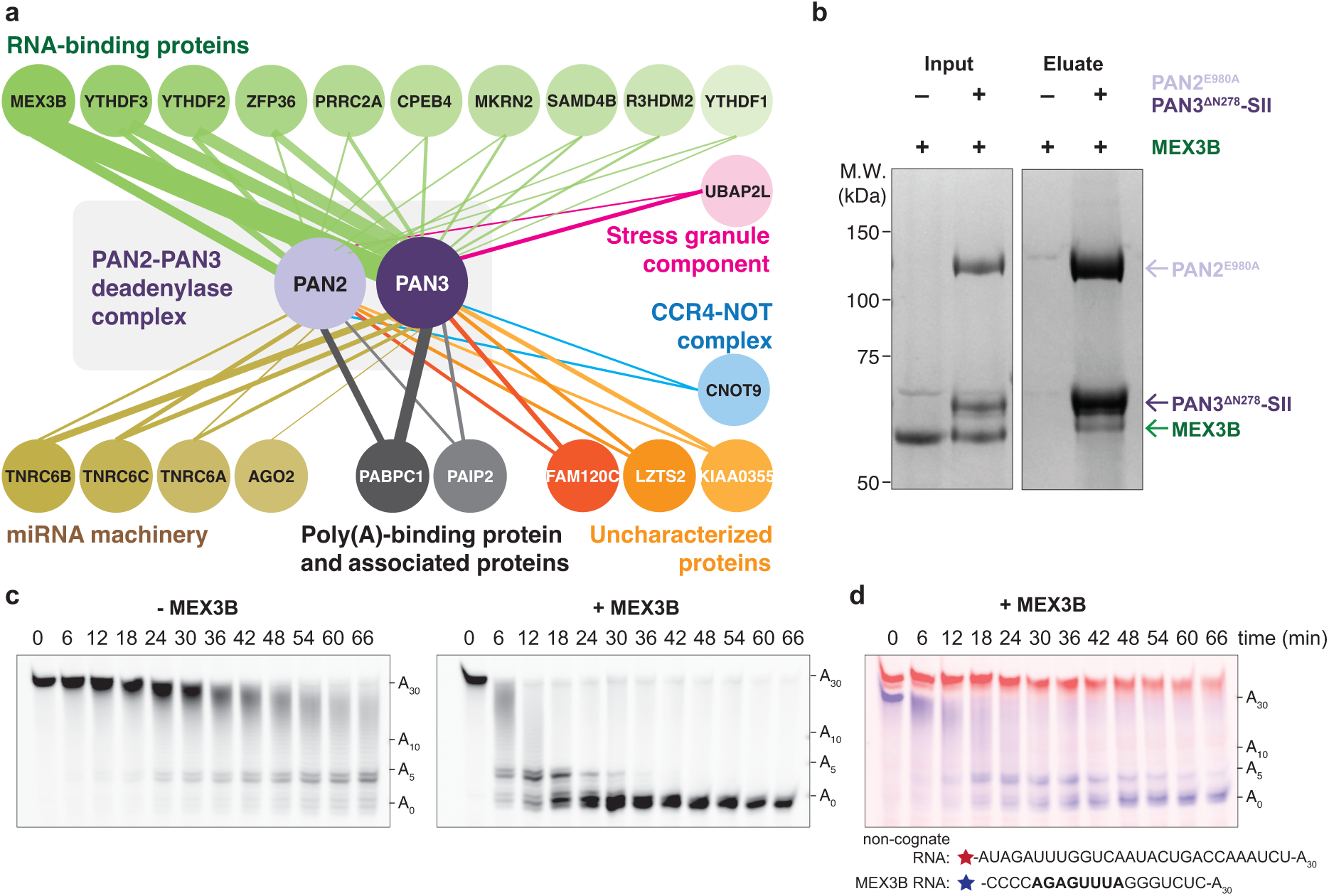
MEX3B interacts with PAN2–PAN3 and mediates targeted deadenylation. (a) **P-body and stress granule components are proximal to PAN2–PAN3 in cells.** PAN2- and PAN3-centric proximity analyses by BioID (Youn et al., 2018) reveal RNA-binding proteins (green), the miRNA machinery (brown), and poly(A)-binding protein (grey) as potential interactors. In addition, a subunit of the CCR4–NOT complex (blue), a stress granule protein (pink), and uncharacterized proteins (orange) are proximal to PAN2– PAN3. AGO2 is proximal to PAN3 and not PAN2 but is shown due to its involvement in miRNA-mediated regulation of gene expression. Each circle represents a prey or BirA*-tagged bait protein, line thicknesses are proportional to fold enrichment of PAN2 and PAN3 over negative controls, and the shade of each circle is colored according to its rank based on average enrichment (darker is more enriched). **(b) MEX3B directly interacts with PAN2–PAN3.** SDS-PAGE analysis of pull-down assays carried out using purified MEX3B and immobilized PAN2^E980A^–PAN3^ΔN278^-SII, or a negative control where no protein was immobilized. Input: immobilized protein + MEX3B in solution; eluate: biotin elution. **(c) MEX3B promotes PAN2–PAN3 deadenylation *in vitro*.** A synthetic RNA substrate was designed to contain a 30-nt poly(A) tail downstream of 19 ribonucleotides containing the binding site for MEX3B (MEX3 Recognition Element, MRE). A MEX3B-RNA complex was formed and deadenylation assays were performed with 40 nM PAN2–PAN3^ΔN278^-SII. Reactions were analyzed by denaturing PAGE. The sizes of RNAs with various poly(A) tail lengths are indicated. **(d) MEX3B mediates selective targeted deadenylation *in vitro*.** Deadenylation of an equimolar mixture of 100 nM fluorescein-labelled MEX3B target RNA (blue) and 100 nM Alexa647-labelled non-cognate RNA (red) containing a 30-nt poly(A) tail with 20 nM PAN2–PAN3^ΔN278-SII^. The MEX3B binding motif is in bold. A 1:1 MEX3B:cognate RNA ratio was used to maximize specific (cognate) interactions over non-specific (non-cognate) interactions.

The identified PAN2–PAN3 interacting RBPs are linked to regulation of specific transcripts in cells. For example, MEX3 is a highly-conserved eukaryotic protein with four paralogs in humans (MEX3A-D) (Jasinski-Bergner et al. 2020) that play roles in diverse physiological processes, including metabolism, the immune response, and development (Chao et al. 2019; Huang et al. 2018; Jiao et al. 2012). MEX3 contains two tandem RNA-binding KH domains that bind to the MEX3 response element (MRE), a bipartite RNA sequence motif (Pagano et al. 2009; Yang et al. 2017). MEX3 represses translation (Draper et al. 1996) and accelerates the deadenylation and decay (Cano et al. 2012, 2015) of MRE-containing transcripts. Second, metazoan YTHDF proteins specifically recognize *N*6-methylated adenosine (m^6^A) modifications via their YTH domains (Luo and Tong 2014; Theler et al. 2014; Xu et al. 2014). Cytoplasmic YTHDF proteins downregulate m^6^A-containing transcripts by recruiting CCR4–NOT through an unknown mechanism (Du et al. 2016; Liu et al. 2020; Wang et al. 2014). Third, ZFP36 (Tristetraprolin or TTP) binds AU-rich elements (AREs) via its zinc finger domains and binds CCR4–NOT to direct rapid deadenylation and degradation of its target transcripts (Fabian et al. 2013; Sandler et al. 2011; Pekovic et al. 2025).

Since the PAN2–PAN3-proximal RBPs identified here are linked to downregulation and deadenylation of their target mRNAs, we hypothesized that they could function as RNA adaptors for PAN2–PAN3.

### MEX3 interacts with PAN2–PAN3 and promotes deadenylation on target transcripts

We next tested if MEX3B directly interacts with PAN2–PAN3. MEX3B was the highest confidence hit that recovered the most peptides for PAN2–PAN3 across all BioID bait proteins. We used purified recombinant MEX3B and a recombinant human PAN2–PAN3 comprised of a catalytically inactive PAN2 (E980A) and StrepII (SII)-tagged PAN3 containing a shortened IDR that improved expression and stability (PAN2^E980A^–PAN3^ΔN278^-SII) (**Figure S1b**). Pull-down assays showed that MEX3B directly interacts with this PAN2– PAN3 construct, in the absence of RNA or other protein factors (**Figure 1b**).

To understand whether MEX3B acts as an RNA adaptor for PAN2–PAN3-mediated deadenylation, we tested its effect in *in vitro* deadenylation assays. We used a synthetic RNA substrate containing a 5′ fluorescent label, followed by a 19-nt sequence containing the MEX3B motif, and a 30-nt poly(A) tail at the 3ʹ end (**Figure S1c**). In the absence of MEX3B, deadenylation by PAN2–PAN3^ΔN278^-SII proceeds relatively slowly under our assay conditions; after 1 h, a substantial fraction of polyadenylated RNA remains (**Figure 1c**).

We next pre-incubated MEX3B with the RNA substrate (**Figure S1c**) and used the resulting RNP complex in deadenylation assays. Strikingly, MEX3B strongly accelerated deadenylation, with fully deadenylated RNA appearing at much earlier time points (6 min) (**Figure 1c**). MEX3B also promotes deadenylation by full-length PAN2–PAN3-SII (**Figure S1d**). Therefore, MEX3B interacts with PAN2–PAN3 and RNA, and increases deadenylation rates *in vitro*.

MEX3B could promote targeted deadenylation by selectively favoring deadenylation of its target RNAs; alternatively, MEX3B could allosterically activate PAN2–PAN3. To differentiate between these two possibilities, we performed two-color competition assays. We mixed fluorescein-labelled MEX3B-target RNA (blue) and Alexa647-labelled non-cognate RNA (red), both with 30-nt poly(A) tails, in equimolar amounts and used them in a deadenylation assay. PAN2–PAN3 deadenylates the MEX3B-target RNA more rapidly than the non-cognate RNA (**Figure 1d**). Therefore, the ability of MEX3B to promote deadenylation is dependent on its interaction with a target RNA, not on allosteric activation of the enzymatic activity.

Together, our data show that MEX3B can interact with PAN2–PAN3 and can promote accelerated deadenylation specifically on its cognate RNA substrate, thereby acting as an RNA adaptor for PAN2–PAN3 deadenylation.

### The C-terminal IDR of MEX3C and PAN3 interact

All four MEX3 paralogs contain two highly-conserved tandem KH domains, N- and C-terminal intrinsically disordered regions (IDRs) and a C-terminal RING domain (**Figure S2a and Supplementary Information 1**). We tested MEX3A, MEX3C and MEX3D and found that they also promote targeted deadenylation by PAN2–PAN3 (**Figure S2b-d**).

MEX3C is the best characterized paralog in terms of its RNA-binding sequence specificity and its physiological role as a post-transcriptional allotype-specific regulator of MHC-I expression (Cano et al. 2012, 2015; Yang et al. 2017). Therefore, we focused on elucidating how MEX3C accelerates deadenylation by PAN2–PAN3. MEX3C and PAN2– PAN3 directly interact in a pull-down assay (**Figure S2e**). We tested how MEX3C influences deadenylation on an RNA substrate containing a 60-nt poly(A) tail bound by PABPC1. MEX3C also accelerates PAN2–PAN3 on PABPC1-bound tails (**Figure S2f**).

Using truncation mutants, we found that multiple regions of MEX3C are involved in the interaction with PAN2–PAN3 and in promoting deadenylation, with the C-terminal IDR (residues 418-600) making a substantial contribution (**Figure S3**). PAN2–PAN3 is comprised of one PAN2 molecule and a PAN3 dimer (**Figure S4a**). We performed pull-down assays with immobilized PAN2–PAN3 and the C-terminal IDR of MEX3C (residues 418-600). These showed that a C-terminal structured region of PAN3, including pseudokinase, coiled-coil, and C-terminal knob domains (termed PKC, residues 463-887), is necessary and sufficient for direct interaction (**Figure S4b-c**). Moreover, deadenylation by PAN2–PAN3^PKC^ was accelerated by MEX3C(222-600), but the catalytic domain of PAN2 was not (**Figure S4d-e**). Notably, deadenylation by PAN2–PAN3^PKC^ with MEX3C(222-600) is slower compared to longer constructs (compare **Figures S3d and S4d**) suggesting that other regions of the proteins also contribute to accelerate PAN2-PAN3 deadenylation.

In summary, multiple regions of PAN2–PAN3 and MEX3C are likely involved in mediating their interaction, consistent with multivalency and similar to interactions between CCR4–NOT and RNA adaptors. We show that PAN3^PKC^ and the MEX3C C-terminal IDR directly interact, and this contributes to targeted poly(A) tail removal by PAN2–PAN3.

### A peptide motif in the C-terminal IDR of MEX3C contributes to binding PAN3

To further analyze how the MEX3C C-terminal IDR interacts with PAN3, we used solution NMR. First, using isotopically labelled MEX3C (residues 418-600, **Figure 2a**), we assigned 94% of the backbone resonances (172/183 residues), including 78% (18/23) of prolines of the C-terminal IDR (**Figure 2b, grey**). The narrow ^1^H dispersion of backbone resonances in the ^1^H, ^15^N HSQC spectrum is a signature for intrinsically disordered proteins. Thus, this region of MEX3C does not have substantial structural order.

**Figure 2.**
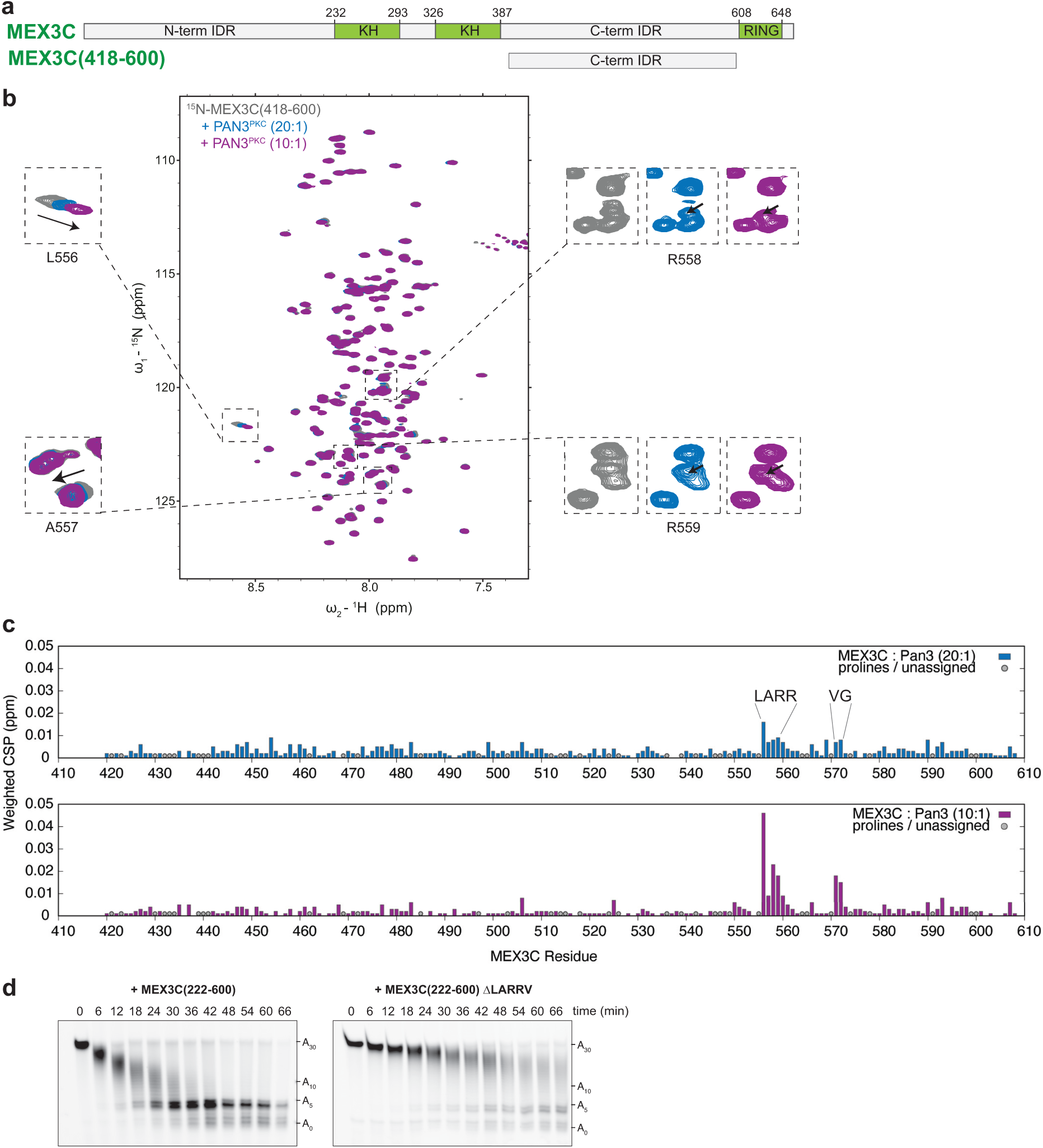
A LARR motif in the C-terminal IDR of MEX3C contributes to increasing the rate of PAN2– PAN3 deadenylation. (a) **Domain diagram of MEX3C and MEX3C(418-600).** MEX3C comprises an N-terminal intrinsically disordered region (IDR), two tandem KH domains which bind a specific RNA sequence, a C-terminal IDR, and a RING domain at the extreme C-terminus. **(b) NMR analysis of MEX3C(418-600).** ^1^H,^15^N 2D HSQC spectra of 45 μM ^15^N-labelled MEX3C(418-600) alone (grey), with 2.25 μM PAN3^PKC^ (blue), or 4.5 μM PAN3^PKC^ (purple). Insets show chemical shift perturbations for resonances of the LARR motif upon addition of PAN3^PKC^ (grey to blue to purple). The arrows indicate the direction of shift upon titration of PAN3^PKC^. **(c) Weighted chemical shift perturbations (CSP) of MEX3C(418-600).** Histogram plots of chemical shift perturbation in ^1^H, ^15^N 2D HSQC upon addition of PAN3^PKC^ at 20:1 (blue, top) and 10:1 (purple, bottom) ratios. Prolines and unassigned residues are indicated with a circle. **(d) The LARR motif contributes to targeted deadenylation by PAN2–PAN3.** Deadenylation assays with 40 nM PAN2–PAN3^ΔN278^-SII were performed with MBP-MEX3C(222-600)-lipoyl or with the same construct containing a deletion of the LARRV sequence. Reactions were analyzed by denaturing PAGE. The sizes of the substrate with various poly(A) tail lengths are indicated.

Next, addition of PAN3^PKC^ to MEX3C(418-600) resulted in chemical shift perturbations in the NMR spectrum in a concentration-dependent manner (**Figure 2b**). The absence of substantial peak broadening upon binding of the 100 kDa PAN3^PKC^ dimer suggests a binding affinity in the high micromolar range (**Figure 2b, insets**). The small number of residues involved further suggests that the interaction with PAN3^PKC^ does not involve extensive conformational changes or long segments of MEX3C. Chemical shift perturbations are observed for resonances corresponding to residues 556-559 which has a sequence of LARR **(Figure 2b-c**), indicating that this may be a short linear motif (SLiM) involved in binding PAN3^PKC^. A VG dipeptide motif downstream of the LARR also shows some perturbation upon addition of PAN3^PKC^. Together, these data suggest that LARR in MEX3C may be involved in binding to PAN3^PKC^.

### The LARR motif contributes to recruitment of PAN2–PAN3 for deadenylation

Unlike most of the C-terminal IDR, the LARR motif is conserved among MEX3 paralogs (**Supplementary Information 1-2**), suggesting that it may be important for MEX3 protein function. Other residues can be found in the second position of this motif, including proline in MEX3A and serine in MEX3D. A small hydrophobic residue (Val, Pro, or Gly) is generally found after the LARR.

To test the importance of the LARR motif in recruiting PAN2–PAN3, we purified MEX3C(222-600) variants where LARRV was mutated. After deletion of LARRV, MEX3C still binds RNA but has impaired ability to promote deadenylation of its cognate RNA by PAN2–PAN3 *in vitro* (**Figure 2d and S5a-b**). Mutation of L556 to aspartic acid or mutation of R558 and R559 to alanine also reduced acceleration of PAN2–PAN3 deadenylation (**Figure S5c**). LARR motifs in the other three MEX3 paralogs are also important for promoting deadenylation (**Figure S5d-h**). Therefore, the conserved LARR motif contributes to targeted deadenylation by PAN2–PAN3.

The importance of the LARR motif raised the possibility that its presence in an IDR of an RBP may be sufficient for targeting of PAN2–PAN3. We introduced the LARRV sequence into the N-terminal IDR of MEX3C(1-422). However, this does not substantially alter PAN2–PAN3 activity (**Figure S5i**). Thus, LARR is necessary, but not sufficient, for full acceleration of PAN2–PAN3-mediated targeted deadenylation.

Overall, these results suggest that the LARR motif contributes to deadenylation. Other portions of the MEX3C are required for full acceleration of PAN2–PAN3 activity, likely in a multivalent mechanism.

### ZFP36, YTHDF2 and YTHDF3 also function as RNA adaptors for PAN2–PAN3

Given the ability of MEX3 proteins to promote targeted deadenylation by PAN2– PAN3, we analyzed other RBPs that were identified as proximal interactors (**Figure 1a**). We tested recombinant ZFP36, YTHDF2 and YTHDF3 and found that all three directly interact with PAN2–PAN3 (**Figure 3a-b**).

**Figure 3.**
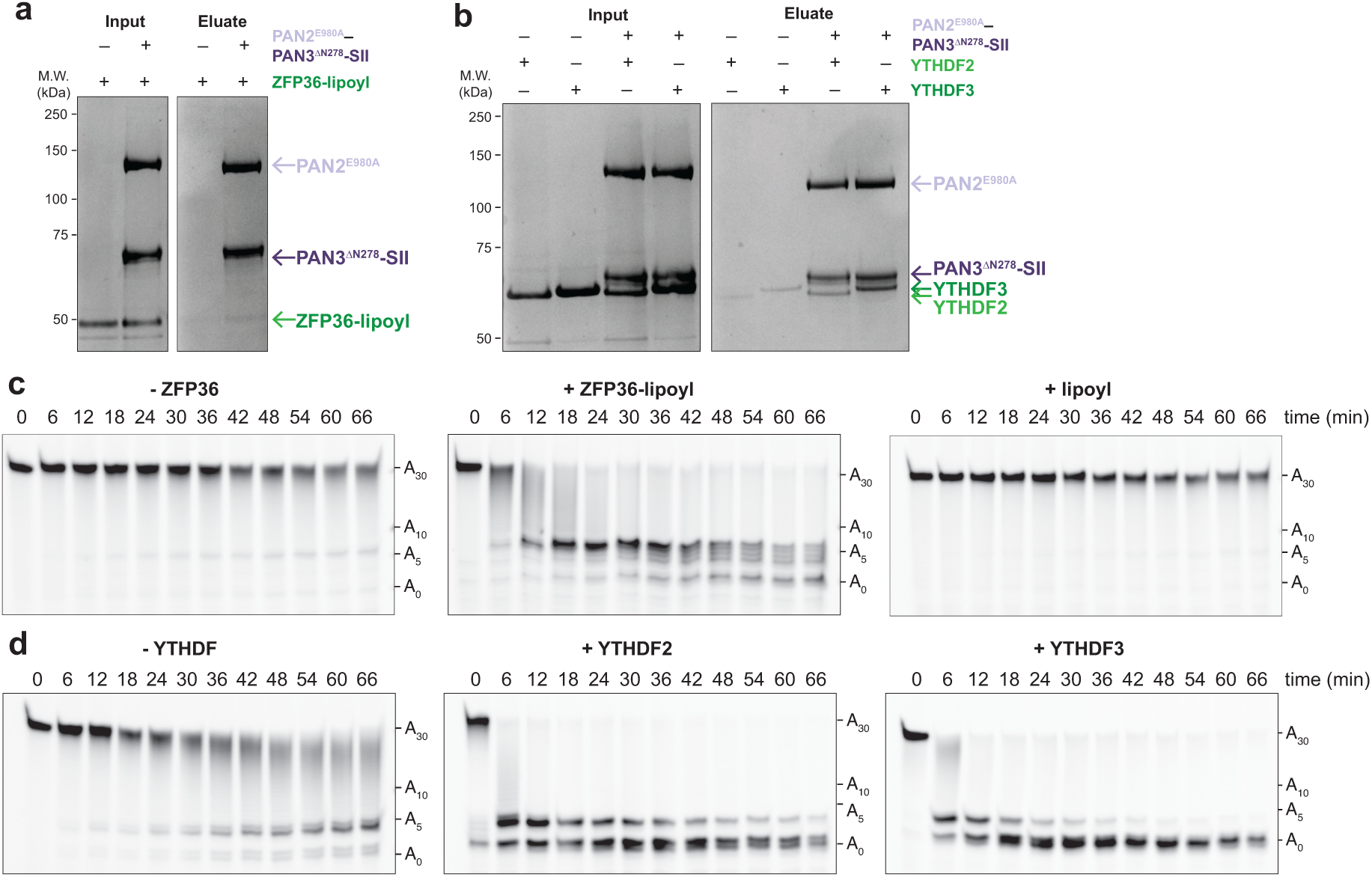
ZFP36, YTHDF2 and YTHDF3 interact with PAN2–PAN3 and mediate targeted deadenylation. (a-b) **ZFP36, YTHDF2 and YTHDF3 directly interact with PAN2–PAN3.** SDS-PAGE analysis of pull-down assays carried out using purified ZFP36-lipoyl (a), YTHDF2 or YTHDF3 (b) and immobilized PAN2^E980A^–PAN3^ΔN278^-SII, or a negative control where no protein was immobilized. Input: immobilized protein + RBP in solution; eluate: biotin elution. **(c-d) RBP interactors promote PAN2–PAN3 deadenylation *in vitro*.** Synthetic RNA substrates were designed to contain a 30-nt poly(A) tail downstream of the binding site for ZFP36 (AU-rich element, ARE) (c) or YTHDF proteins (consensus m6A-containing motif) (d). RBP-RNA complexes were formed and deadenylation assays were performed with 40 nM PAN2–PAN3^ΔN278^-SII. The lipoyl domain alone was included as a negative control in panel (c). Reactions were analyzed by denaturing PAGE. The sizes of RNAs with various poly(A) tail lengths are indicated.

To test if these interactions can promote targeted deadenylation, we designed fluorescently-labelled synthetic RNAs containing the binding motifs for these RBPs (m^6^A or AU-rich element, ARE) as well as a 30-nt 3ʹ poly(A) tail. Using our *in vitro* deadenylation assay, we found that all three of these RBPs promote PAN2–PAN3-mediated deadenylation on their target mRNAs, but not on non-target mRNAs (**Figure 3c-d and S6**).

Together, this shows that multiple RBPs can interact with PAN2–PAN3 and can promote accelerated deadenylation specifically on their cognate RNA substrates, thereby acting as RNA adaptors for PAN2–PAN3 deadenylation.

### A degenerate LARR motif is found in ZFP36 and YTHDF3

The contribution of the LARR motif in MEX3 proteins led us to search for similar motifs in the ZFP36 and YTHDF proteins. Human ZFP36 contains an LARR motif (residues 240-244) in a C-terminal IDR (**Figure 4a**) that is partially conserved in mice (VTRR), but not in *Xenopus* (**Supplementary Information 3**). To test the importance of this motif in promoting deadenylation by PAN2–PAN3, we deleted LARRD from human ZFP36. This impaired (but did not eliminate) its ability to increase the rate of PAN2–PAN3 deadenylation (**Figure 4b and S7a-b**).

**Figure 4.**
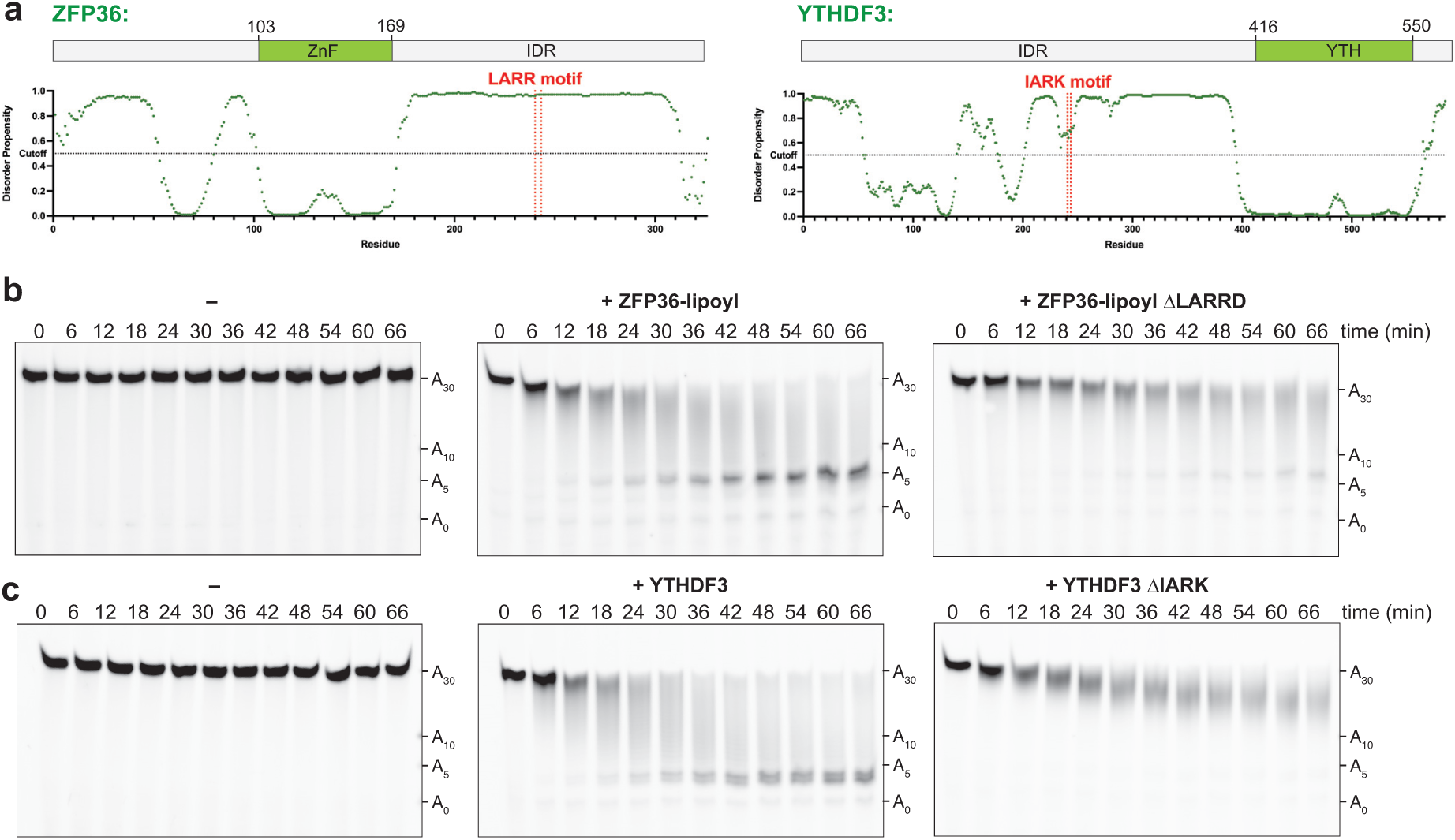
A degenerate LARR motif contributes to RBP-mediated targeted deadenylation. (a) The domain organization and disorder prediction of ZFP36 and YTHDF3. ZFP36 contains N- and C-terminal IDRs and a central RNA-binding domain comprising tandem zinc fingers. An LARR motif (annotated in red) is found in the C-terminal IDR.YTHDF3 contains an extensive, mostly disordered, N-terminal region and a C-terminal YTH RNA-binding domain. An IARK motif (annotated in red) is found in the N-terminal IDR. **(b) The LARRD motif of ZFP36 contributes to acceleration of PAN2–PAN3 deadenylation.** Wild-type ZFP36 or a construct lacking the LARRD motif, both with C-terminal lipoyl solubility tag, were added to the ARE-containing RNA with a 30-adenosine tail, prior to adding 20 nM PAN2–PAN3^Δ278^-SII. Reactions were analyzed by denaturing PAGE. The sizes of the substrate with various poly(A) tail lengths are indicated. **(d) The IARK motif of YTHDF3 contributes to acceleration of PAN2–PAN3 deadenylation.** Wild-type YTHDF3 or a construct lacking the IARK motif was added to a GGm^6^ACU-motif-containing RNA with a 30-adenosine tail, prior to adding 5 nM PAN2–PAN3^Δ278^-SII. Reactions were analyzed by denaturing PAGE. Substrate sizes with various poly(A) tail lengths are indicated.

YTHDF3 contains an IARK motif (residues 241-244) in its extensive N-terminal IDR (**Figure 4a**). The highly related YTHDF1 and YTHDF2 proteins contain an IASK motif, and YTHDC1, a nuclear YTH domain-containing protein which is not expected to be involved in cytoplasmic deadenylation, does not harbor a recognizable motif (**Supplementary Information 4**). The IARK motif is conserved in YTHDF3 vertebrate homologs (**Supplementary Information 5**). We purified a construct of YTHDF3 lacking IARK and found that this impaired, but did not eliminate, the ability of YTHDF3 to accelerate PAN2– PAN3 deadenylation on its cognate RNA (**Figure 4c and S7c-d**).

Taken together, these data suggest that LARR-like motifs can play a role in some RBPs to facilitate targeted deadenylation of specific transcripts by PAN2–PAN3. In all examined cases, this motif occurs in intrinsically disordered regions, suggesting that flexibility surrounding the motif may be required for its function. Finally, we note that all proteins which had deletions of the LARR-like motif retained an ability to promote PAN2– PAN3-mediated deadenylation, suggesting that additional regions of these RBPs are likely to contribute. This is consistent with RNA adaptors interacting with PAN2–PAN3 via a multivalent mechanism, similar to how RNA adaptors interact with CCR4–NOT.

### PAN2–PAN3 is proximal to additional RBPs

All four of the RBPs we tested (MEX3B, ZFP36, YTHDF2 and YTHDF3) act as RNA adaptors for PAN2–PAN3 *in vitro*. To understand whether further RBPs may be involved in PAN2–PAN3 targeted deadenylation, we performed proximity-based proteomics on PAN2 and PAN3. We fused two alternative biotin ligases (BirA*-FLAG or miniTurbo-FLAG) to the N- or C-termini of PAN2, PAN3, and PAN3^PKC^ and performed proximity-dependent biotinylation analyses in HEK293 cells (**Figure S8a and Table S2**).

We merged the resultant data from N- and C-terminal tags and then compared the miniTurbo and BirA* labelling datasets for each of PAN2, PAN3, and PAN3^PKC^ (**Figure S8b-c, Table S3**). This revealed known interactors of the PAN2–PAN3 complex, including TNRC6 proteins and PABPC1 as well as the MEX3 and YTHDF proteins. PAN2 had the fewest high-confidence preys common to both labelling techniques, suggesting that at least one of the tags interfered with protein localization or folding. In contrast, the dataset from tagged full-length PAN3 resulted in many common high-confidence proximity interactors (**Figure S8c**). PAN3^PKC^ had an intermediate number of high-confidence preys.

70 high-confidence prey proteins were identified for full-length PAN3. We used dot plot analysis to compare the interactors across datasets (**Figure 5a**, shown for n=55 proteins common to all four PAN3 datasets), and annotated them by function according to the Gene Ontology Project to select RNA-binding proteins (Baltz et al. 2012; Castello et al. 2012; Gaudet et al. 2011). In total, we identified 46 cytoplasmic RBPs (including the MEX3 and YTHDF proteins), which may act as RNA adaptor proteins for PAN2-PAN3.

**Figure 5.**
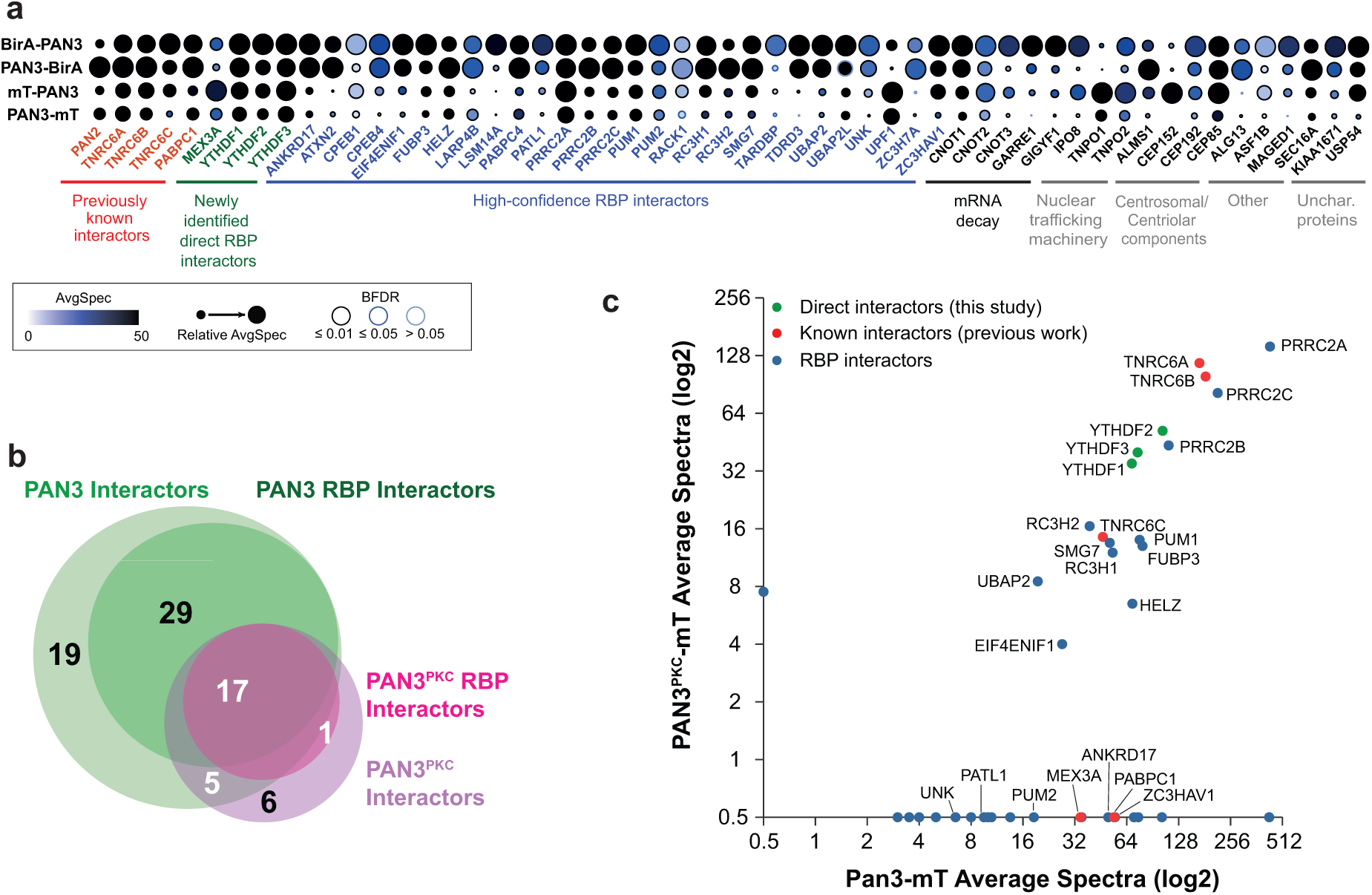
Identification of additional RNA-binding protein interactors of PAN2–PAN3 by proximity labelling. (a) **Annotated dot plot of high-confidence interactors of PAN3.** PAN3 was tagged at the N- and C-termini by BirA* (BirA_PAN3, PAN3_BirA) or miniTurbo (mT_PAN3, PAN3_mT) and used in proximity labelling assays (rows). Selected prey proteins were annotated by color for function as indicated. The dotplot is annotated by averaged spectral count (light blue to black), relative averaged spectral count between groups (dot size), and Bayesian False Discovery Rate (BFDR; outer circle color). **(b) The RBP proximity interactomes of PAN3^PKC^ and PAN3 largely overlap.** Spectra were averaged for N- and C-terminally tagged versions of PAN3 and PAN3^PKC^. Prey proteins which were identified with both BirA* and miniTurbo tags are shown (PAN3: light green, n=70, PAN3^PKC^: light magenta, n=29). These high-confidence proximity interactors were filtered by known RNA-binding function (PAN3, n=46; PAN3^PKC^, n=18). Most RBP interactors of PAN3^PKC^ (17/18) are also proximal to PAN3. **(c) The proximity interactome of full-length PAN3 recapitulates most interactors of PAN3^PKC^.** Data from Figure 5b were used to generate a bait-bait scatter plot comparing proximal RBP interactors of PAN3 and PAN3^PKC^; each dot represents a high-confidence proximal prey RBP (BFDR ≤ 0.01) and selected dots are labelled. 17 high-confidence RBP interactors of both PAN3 and PAN3^PKC^ lie within the graph body. Red: previously known interactors of PAN2–PAN3; green: direct interactors of PAN2–PAN3 characterized in this study; blue: putative proximal RBP interactors of PAN2–PAN3.

Since the PKC domain of PAN3 interacts directly with the C-terminal IDR of MEX3C, it may act as a hub for protein-protein interactions. We thus repeated the dot plot analysis for PAN3^PKC^ as above (n=44 in at least three PAN3^PKC^ datasets) and identified 18 cytoplasmic RBPs as high-confidence proximity interactors (**Figure S8d**). Strikingly, there was a large overlap between prey RBPs for PAN3^PKC^ and PAN3 (**Figure 5b**). These include known interactors (**Figure 5c; red, green**) and novel, putative RNA-binding protein interactors (**Figure 5c, blue**). Since proximity-based proteomics of PAN3^PKC^ recapitulates a large number of interactors identified with full-length PAN3, the PKC region of PAN3 may act as a hub of protein-protein interactions.

Overall, our proximity-based proteomics study identifies a number of potential RNA adaptors for PAN2–PAN3. For example, the PUM protein family (PUM1, PUM2) and the Roquin family (RC3H1, RC3H2) both contain extensive intrinsically disordered regions and RNA-binding domains, which recognize specific RNA sequence elements. Both protein families regulate transcript half-lives by recognizing specific RNA sequence motifs and promoting target mRNA degradation in response to specific cellular cues (Goldstrohm et al. 2018; Schaefer and Klein 2016), and therefore may represent additional RNA adaptor proteins for PAN2–PAN3.

Interestingly, several high-confidence proximal interactors of the PAN2–PAN3 complex also act as mRNA adaptor proteins for the CCR4–NOT deadenylase complex. This raised the possibility that PAN2–PAN3 and CCR4–NOT could interact with the same mRNA adaptor proteins to promote targeted deadenylation on common mRNAs. We compared high-confidence proximal prey proteins (n=46) from the BirA*-tagged PAN3 dataset to a filtered list of proximal RBP interactors from a previous dataset of BirA*-tagged CNOT9 (n=49), a constitutive component of CCR4–NOT (Youn et al. 2018) (**Figure 6a and Table S3**). These studies were carried out in the same cell line under similar experimental conditions, facilitating a direct comparison. Surprisingly, the majority of RBP interactors of PAN3 (green; 33 of 46) is also proximal to CNOT9 (brown), and vice versa (33 of 49), including TNRC6, YTHDF, Roquin, and PUM proteins (**Figure 6b**). The large overlap in high-confidence prey interactors of both deadenylase complexes suggests that they may share RNA adaptor proteins that mediate deadenylation specificity.

**Figure 6.**
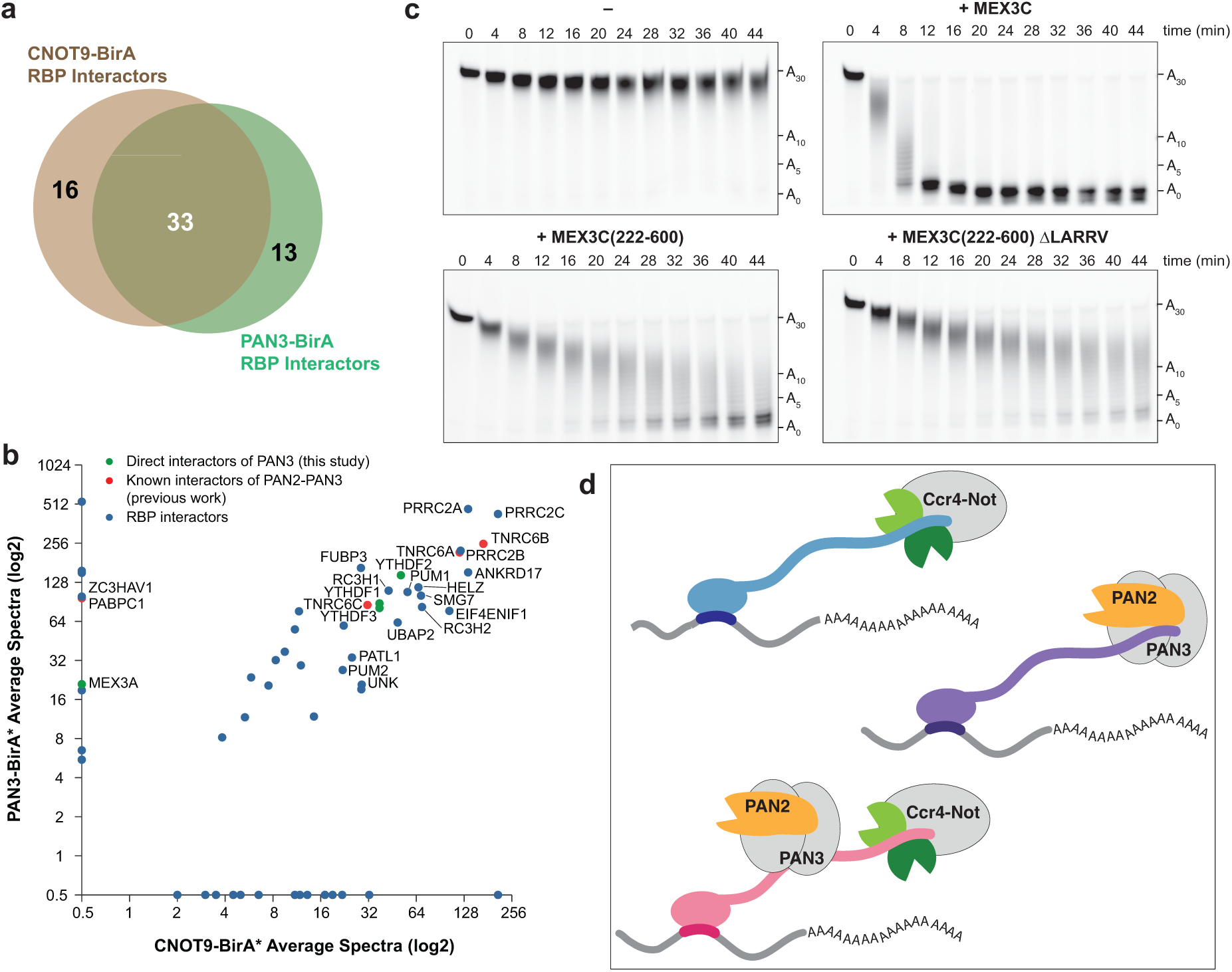
RNA adapters for PAN2–PAN3 and CCR4–NOT are highly redundant. (a) **Many RBP interactors of PAN3 are also proximal to CCR4–NOT.** Prey RBPs from N- and C-terminally averaged BirA*-tagged CNOT9 (n=49) were compared to high-confidence prey RBPs from N- and C-terminally averaged BirA*-tagged PAN3 (green, n=46). Many RBP interactors of PAN3 (33/46) were also proximal to CNOT9. **(b) Sequence-specific RBPs are proximal to PAN2–PAN3 and CCR4–NOT.** Data from Figure 6a were used to generate a bait-bait scatter plot comparing RBP proximal interactors of PAN3 and of CNOT9; each dot represents a high-confidence proximal prey RBP (BFDR ≤ 0.01) and selected dots are labelled. Red: previously known interactors of PAN2–PAN3; green: direct interactors of PAN2–PAN3 characterized in this study; blue: proximal RBP interactors of PAN3. **(c) MEX3C accelerates CCR4–NOT deadenylation *in vitro.*** Deadenylation assays were performed with 10 nM human CCR4–NOT with synthetic RNA substrates containing a 30-adenosine poly(A) tail downstream of 19 non-poly(A) ribonucleotides containing the MEX3 Recognition Element (MRE). Full-length MEX3C, or the C-terminal half of MEX3C without the RING domain (222-600) containing or lacking (ΔLARRV) the LARR motif were added at saturating RBP:RNA ratios. Reactions were analyzed by denaturing PAGE. RNA sizes with various poly(A) tail lengths are indicated. **(d) Tethering model of deadenylation specificity.** RNA-binding proteins (blue, purple, pink) can act as RNA adapters for PAN2–PAN3 and CCR4–NOT. These RNA adapters bind specific sequence motifs to effectively tether the deadenylation complexes to their substrates. PAN2–PAN3 and CCR4–NOT may target different or the same mRNAs for specific deadenylation via unique mechanisms.

ZFP36 and YTHDF proteins are known to act as RNA adaptor proteins for CCR4– NOT, but the effect of MEX3 on CCR4-NOT activity is unknown. Given the high overlap in RBPs that are proximal to PAN2–PAN3 and CCR4–NOT, we tested whether MEX3C directly recruits CCR4–NOT for targeted deadenylation *in vitro*. These assays showed that MEX3C accelerates deadenylation of MRE-containing RNA by CCR4–NOT (**Figure 6c**). Interestingly, in contrast with PAN2-PAN3, MEX3 tethering of CCR4-NOT is more substantially reduced by deletion of the MEX3 N-terminal IDR and is less dependent on the MEX3C LARR motif (compare to **Figure 2d**). Thus, CCR4–NOT and PAN2–PAN3 may be dependent on different parts of MEX3C for deadenylation. Since the two deadenylation complexes require distinct regions of RBPs for targeting, this raises the possibility that they could be simultaneously recruited to the same mRNA for robust downregulation (**Figure 6d**).

## Discussion

Previously, PAN2–PAN3 was thought to be recruited to mRNAs primarily by PABPC1 and thus, to predominantly play a more general role in shortening long poly(A) tails (Schäfer et al. 2019; Yamashita et al. 2005). Here, we identify sequence-specific human RBPs that can directly bind PAN2–PAN3 and recruit it to their target RNAs. By accelerating deadenylation of their cognate transcripts, these RBPs could provide the specificity critical for precisely regulating mRNA stability. Furthermore, in proximal labeling studies, we find a large overlap in RBPs that bind PAN2–PAN3 and CCR4–NOT. The recruitment of both major deadenylases by common RBPs would provide robustness in targeted mRNA degradation.

### A tethering model of targeted deadenylation

RBPs are frequently modular, containing RNA-binding domains and extensive IDRs (Jonas and Izaurralde 2013; Zhao et al. 2021). In the examples studied here, both the RNA binding-domain and a PAN3-interacting IDR are required for robust acceleration of PAN2– PAN3 deadenylation. We therefore propose that RBPs act as flexible tethers between PAN2– PAN3 and specific substrate mRNAs to promote rapid deadenylation and downstream decay (**Figure 6d**). Specificity of binding of the RBP to RNA can be achieved via the mRNA sequence alone or via RNA modifications (e.g. in the case of YTH proteins). Overall, this suggests that the mechanism of specificity in deadenylation is common to both PAN2–PAN3 and CCR4–NOT.

IDRs often exhibit high evolutionary plasticity, as they can arise *de novo* or evolve convergently due to relatively low selection pressures (Tompa et al. 2014). We found that some vertebrate PAN2–PAN3 RNA adaptor proteins contain a LARR-like motif that contributes to tethering. Interactions between short LARR-like sequences and PAN3 would likely be low-affinity and transient. Significantly, we found that the LARR motif alone was not sufficient to mediate targeted deadenylation; sequences outside the LARR must also contribute to binding. Multivalency is therefore likely important for interaction of RNA adaptors with both CCR4–NOT (Stowell et al. 2024; Pekovic et al. 2025) and PAN2–PAN3.

CCR4–NOT also interacts with SLiMs found in IDRs of RNA adaptor proteins. Some of these motifs form an ɑ helix upon interaction (Fabian et al. 2013; Raisch et al. 2016), and can involve large regions of the IDR (Stowell et al. 2024; Pekovic et al. 2025, Webster et al. 2019). Interestingly, in the case of MEX3, CCR4–NOT and PAN2–PAN3 do not appear to have overlapping binding sites on RNA adaptor proteins. Thus, some RNA adaptor proteins may flexibly tether both deadenylation complexes to specific mRNAs simultaneously, increasing their local concentration and thereby promoting rapid mRNA decay (**Figure 6d**).

### PAN3 is a hub for protein-protein interactions

The majority of previously identified interactors of PAN2–PAN3, such as the poly(A)-binding protein PABPC1 or TNRC6 (GW182) proteins of the miRNA machinery, are thought to interact with the PAN3 subunit: PABPC1 binds the conserved PAM2 peptide motif in the PAN3 N-terminal IDR (Kozlov et al. 2010; Mangus et al. 2004) whereas tryptophan residues of GW182 interact with a Trp-binding pocket on the PKC dimer of PAN3 (Braun et al. 2011; Christie et al. 2013; Huntzinger et al. 2013; Kuzuoglu-Öztürk et al. 2012; Schäfer et al. 2019). We show here that the C-terminal dimeric PKC region of PAN3 is important for the direct interaction with MEX3, and likely other RBPs. Taken together, our data provide a molecular explanation for previous suggestions that PAN3 acts as a regulatory subunit of PAN2–PAN3 (Brown et al. 1996; Uchida et al. 2004) by acting as a hub for protein-protein interactions that regulate deadenylation.

It remains unclear whether different (or additional) sequence motifs in other RBPs also promote interaction with PAN2–PAN3 and whether interactions with RBPs are mutually exclusive, such that only one RBP can interact with PAN2–PAN3 at a time. In certain cases, multiple cis-acting 3ʹ-UTR elements binding to different RBPs may act synergistically to recruit PAN2–PAN3 to facilitate specific degradation (Plass et al. 2017).

### Specificity and redundancy in deadenylation

Using proximity labelling techniques, we find that a large number of RBPs are proximal to PAN2–PAN3; these may regulate deadenylation by acting as RNA adaptors. Some of these putative interactors, including the PUM and RC3H/Roquin families, are known to accelerate mRNA degradation and downregulate gene expression *in vivo* (Essig et al. 2018; Goldstrohm et al. 2018), consistent with roles in targeted deadenylation. The number of interactors of PAN2–PAN3 is likely much higher than what we report here. More extensive screening, for example in different cell lines and cellular conditions, would therefore reveal additional RNA adaptors that provide specificity to PAN2–PAN3 and CCR4-NOT. The dynamic interactome of deadenylation complexes would enable a more complete understanding of specificity in deadenylation and post-transcriptional regulation of gene expression.

Intriguingly, our study identified a substantial overlap between RBPs that are proximity interactors of both PAN2–PAN3 and CCR4–NOT. This suggests that there is substantial functional redundancy and interplay between the two major deadenylase complexes. This is consistent with previous data demonstrating that deletion of an individual deadenylase enzyme in yeast had a small effect on global poly(A) tail lengths, whereas simultaneous deletion of both Pan2 and Ccr4 resulted in growth defects and almost no observable deadenylation (Boeck et al. 1996; Tucker et al. 2001, 2002). Potential redundancy of deadenylation enzymes (CNOT6, CNOT6L, CNOT7, CNOT8, PAN2 in humans) and the essentiality of mRNA decay complicates the study of this process in mammalian cells, explaining why the role of PAN2–PAN3 in targeted deadenylation had not been recognized previously.

Upon stimulation by specific cellular cues, it may be advantageous for targeted RNAs to be rapidly degraded, which would be facilitated by the recruitment of both PAN2–PAN3 and CCR4–NOT by the same RBPs to reinforce the downregulated fate of the mRNA (**Figure 6d**). It is possible that the two deadenylation complexes tend to function in different contexts, in different cellular compartments, or on different lengths of poly(A) tails. A recent study suggested that YTHDF3 preferentially recruits PAN2–PAN3, whereas YTHDF2 binds CCR4–NOT, in redundant pathways to mediate m^6^A-targeted transcript degradation (Liu et al. 2020). Thus, certain cellular cues may result in the differential recruitment of either deadenylase. Targeted mRNA decay through parallel deadenylation pathways could therefore further increase the tunability of mRNA degradation.

## Conclusion

A major question in understanding post-transcriptional regulation of gene expression remains how transcripts can be specifically and rapidly targeted for degradation upon changes in cellular conditions. We have identified that PAN2–PAN3 can mediate targeted deadenylation by interacting with numerous RBPs. Targeted deadenylation by PAN2–PAN3 may occur redundantly with that by CCR4–NOT, which would both reinforce the robustness of mRNA degradation, and further increase the complexity of regulation of post-transcriptional gene expression.

## Acknowledgements

We thank Brian Carrick and Megan Gough as well as other members of the Passmore lab for advice and feedback, and J.G. Shi (LMB baculovirus) for support. This work was supported by a Herchel Smith PhD Studentship from the University of Cambridge (to T.T.L.T.); the Medical Research Council, as part of United Kingdom Research and Innovation, MRC file reference number MC_U105192715 (L.A.P.); the European Union’s Horizon 2020 research and innovation programme (ERC Consolidator grant agreement 725685) (to L.A.P); the National Sciences and Engineering Research Council of Canada, RGPIN-2019-06297 (to A.-C.G.); and the Canada Foundation for Innovation and the Ontario Research Fund (to A.-C.G.). A.-C.G. is the Canada Research Chair (Tier 1) in Functional Proteomics.

## Author contributions

T.T.L.T. led the project. T.T.L.T, F.Z. and C.W.H.Y. designed and performed experiments, analyzed the data and wrote the manuscript. J.A.W.S. performed some of the deadenylation assays; A.R. performed initial experiments on MEX3. C.W.H.Y. and S.M.V.F. performed NMR experiments. F.Z. and A.-C.G. performed proximity-based proteomics. L.A.P. and A.-C.G. supervised the project, analyzed the data and wrote the manuscript.

## Rights Retention Statement

This work was supported by the Medical Research Council, as part of United Kingdom Research and Innovation (also known as UK Research and Innovation) [MRC file reference number MC_U105192715]. For the purpose of open access, the MRC Laboratory of Molecular Biology has applied a CC BY public copyright licence to any Author Accepted Manuscript version arising.

## Materials and Methods

### DNA constructs

Full-length *H. sapiens* PAN2, PAN3-SII (SII, twin StrepII), MEX3A-D, YTHDF3, ZFP36, PABPC1, and *H. sapiens* CCR4-NOT subunits (C-terminal SII tag on CNOT8 subunit) were codon-optimized for *E. coli* expression, synthesized, and cloned into pACEBac1 (Epoch Life Science). *H. sapiens* YTHDF2 were synthesized and cloned into pMK by GeneArt (ThermoFisher Scientific). All cloning steps were carried out using Gibson Assembly and plasmids were propagated in TOP10 *E. coli* (Invitrogen).

PAN2–PAN3 complexes (PAN2-GST–PAN3-SII, PAN2–PAN3^ΔN278^-SII, PAN2^E980A^–PAN3^ΔN278^-SII, PAN2–PAN3^PKC^-SII (ΔN460)) were cloned into a modified pBIG1a by Gibson assembly (Hill et al. 2019; Weissmann et al. 2016). Individual PAN2 or PAN3 constructs (PAN2^UCH-Exo^, PAN2^WD40^, PAN3^PKC^) were cloned into pACEBac1 with a C-terminal SII tag. MEX3A-D constructs containing the RNA-binding KH domain were cloned into a pMAL-c5x vector with an N-terminal maltose binding protein (MBP) tag and modified to contain a 3C protease cleavage site. For some MEX3 constructs, a 3C-cleavable C-terminal lipoyl domain from *Bacillus stearothermophilus* pyruvate dehydrogenase was included where indicated for protein stability and solubility (Brandt et al. 2012; Hipps et al. 1994). MEX3C(418-600) constructs and ZFP36 were cloned into a modified pFX vector (Geertsma and Dutzler 2011) containing an N-terminal His_6_-SUMO tag (cleavable by SUMO protease) and a C-terminal 3C-cleavable lipoyl domain. YTHDF constructs were cloned into the modified pMAL-c5x vector as above with a C-terminal SII tag. PABPC1 was cloned into a modified pET28a vector containing a TEV-cleavable C-terminal SII tag. CCR4-NOT subunits were combined for co-expression into a modified pBIG2 by Gibson assembly.

For BioID studies, full-length PAN2, PAN3 and PAN3^PKC^ were amplified with DNA ends compatible with Gateway cloning and cloned into pDONR223 entry vectors. These were used to generate pcDNA-DEST40 vectors by Gateway cloning (ThermoFisher Scientific) containing the bait with N- or C-terminal BirA*-FLAG or miniTurbo-FLAG tags.

### Baculovirus-mediated expression and purification of PAN2–PAN3

pACEBac1 or pBIG1a plasmids encoding the protein or protein complex of interest were transformed into *E. coli* DH10 EMBacY cells (Geneva Biotech), and colonies with successful integration were identified by blue/white selection and SwaI digestion. Bacmid DNA was isolated using previous protocols (Stowell et al. 2016). *Sf*9 insect cells were grown in Insect-XPRESS medium (Lonza) at 27°C. 10 µg bacmid DNA was transfected into 1×10^6^ adherent *Sf*9 cells using 20 µl FuGENE (Promega) and allowed to incubate for 72h. YFP-positive supernatants were filter-sterilized and stored with 50% FBS. Primary virus was amplified by infecting 50 ml suspension *Sf*9 cells at 2×10^6^ cells/ml; cells were monitored for viability, fluorescence, and density every 24 h. Secondary virus was harvested by filter-sterilization, and protein expression was monitored by a small-scale pull-down on the tag of interest. For large-scale protein expression, 4.5 ml secondary virus was added to 500 ml *Sf*9 cells in suspension at 1.5×10^6^ cells/ml in 2 liter roller bottles. Cells were harvested and centrifuged at 3,876 × g, and flash frozen until use.

All purification steps were performed at 4°C in an AKTA Pure 25 (Cytiva); sample purity and homogeneity were monitored at each stage by SDS PAGE.

For PAN2-GST–PAN3-SII, frozen *Sf*9 pellets were resuspended in 2-3× Buffer A1 (50 mM PIPES pH 6.5, 500 mM NaCl, 2 mM DTT, 0.5 mM PMSF, 1× EDTA-free complete protease inhibitor (Roche), 20 µg/ml DNase I) by volume and lysed by sonication (3s on, 6s off, 3× 1.5 min, 70% amplitude) using a 10-mm tip on a VC750 ultrasonic processor (Sonics). Lysate was centrifuged at 235,418 × g for 30 min, 4°C. Clarified supernatant was incubated with 4 ml equilibrated glutathione sepharose 4B resin (Cytiva) for 2 h, 4°C. Resin was washed with 50 ml Buffer A1, 50 ml Buffer B1 (50 mM PIPES pH 6.5, 1 M NaCl, 2 mM DTT) and protein was eluted with 30 ml Buffer C1 (50 mM PIPES pH 6.5, 300 mM NaCl, 2 mM DTT, 20 mM reduced glutathione). Eluate was pooled and diluted 5× with Buffer D1 (25 mM PIPES pH 6.5, 300 mM NaCl, 1 mM DTT) and loaded onto a 5 ml HiTrap StrepTrap HP column (Cytiva) equilibrated in Buffer D1. The column was washed with 5 CV Buffer D1 and eluted with 5 CV Buffer E1 (Buffer D1 + 20 mM DSB). Fractions containing PAN2-GST–PAN3-SII were pooled and concentrated to ∼5 µM, flash frozen, and stored at -80°C.

For SII-tagged PAN2–PAN3 (PAN2–PAN3^ΔN278^-SII, PAN2^E980A^–PAN3^ΔN278^-SII, PAN2–PAN3^PKC^-SII, PAN2^UCH-Exo^-SII, PAN2^WD40^-SII), frozen *Sf*9 pellets were resuspended in 2-3× Buffer A2 by volume (50 mM HEPES pH 8.0, 500 mM NaCl, 2 mM DTT, 0.5 mM PMSF, 1× EDTA-free cOmplete protease inhibitor (Roche), 20 µg/ml DNase I, 2 ml BioLock (IBA)) and lysed and centrifuged as above. Lysate was incubated with 4 ml StrepTactin resin (IBA) for 2 h, 4°C. Resin was washed with 50 ml Buffer A2, 50 ml Buffer B2 (50 mM HEPES pH 8.0, 1 M NaCl, 2 mM DTT) and protein was eluted with 30 ml Buffer C2 (50 mM HEPES pH 8.0, 200 mM NaCl, 2 mM DTT, 20 mM desthiobiotin). Eluate was pooled and diluted 5× with Buffer D2 (25 mM HEPES pH 8.0, 1 mM DTT) and loaded onto a 5 ml HiTrap Q HP column (Cytiva) equilibrated in Buffer D2 + 50 mM NaCl. The column was washed with 5 CV Buffer D2 + 100 mM NaCl and eluted with a 20 CV 10-50% Buffer E2 gradient (Buffer D2 + 1 M NaCl). Peak fractions were pooled, concentrated, and loaded onto an S200 26/60 column (Cytiva), equilibrated in Buffer F2 (20 mM HEPES pH 8.0, 200 mM NaCl, 1 mM TCEP). Peak fractions were pooled, concentrated to ∼20 µM, flash frozen, and stored at -80°C.

### Baculovirus-mediated expression and purification of CCR4-NOT

Expression and purification of human CCR4-NOT was carried out according to Absmeier, Chandrasekaran and colleagues (Absmeier et al. 2023). In brief, a low multiplicity of infection (MOI) strategy was used to generate 6 liters of *Sf*9 expression culture obtained by dilution of a P0-infected *Sf*9 culture to maintain cell density between 1.5-3.0×10^6^ cells/ml. Cells were harvested as above.

Purification of human CCR4-NOT was carried out as previously described by sequential affinity and ion exchange purification steps (Absmeier et al. 2023). SII-tagged CCR4-NOT complex was isolated by batch-binding of clarified lysate to StrepTactin resin (IBA), followed by further purification on a 5 mL HiTrap Q HP column (Cytiva). Fractions containing the purified complex were pooled and concentrated on a 1 ml Resource Q column. The purified complex was pooled, flash frozen, and stored at -80°C.

### *E. coli* expression and purification of MEX3, YTHDF, ZFP36, and PABPC1

*E. coli* was grown in Terrific Broth (TB) containing the appropriate antibiotic (Ampicillin for pMAL-c5x; Kanamycin for pFX). Modified pMAL-c5x or pFX plasmids encoding the protein of interest were transformed into *E. coli* BL21 DE3 (STAR) (ThermoFisher Scientific) by heat shock. Colonies which expressed the protein of interest were used to inoculate a small-scale overnight culture (∼150 ml). 10 ml of culture was transferred to 1 liter media to initiate growth. Protein expression was induced with 1 mM IPTG when OD_600_ = 0.5-1.0, for 5 h at 37°C. Cells were harvested by centrifugation at 3,876 × g, 20 min and flash frozen until use.

Isotopically labelled proteins were overexpressed in M9 media, supplemented with 1.7g/l yeast nitrogen base lacking NH_4_Cl and amino acids (Merck). We supplemented 1 g/l ^15^NH_4_Cl and 4 g/l ^13^C-glucose for ^15^N and ^13^C labelling, respectively. All cultures were grown in 2 liter conical flasks at 180 rpm, 37°C.

Pellets expressing MBP-tagged proteins (MEX3, YTHDF) were resuspended in 2-3× Buffer A2 by volume, and lysed and centrifuged as above. Lysate was incubated with 4 ml equilibrated amylose sepharose resin (NEB) for 2 h, 4°C. Resin was washed with 50 ml Buffer A2, 50 ml Buffer B2 and protein was eluted in six steps with 30 ml Buffer C3 (50 mM HEPES pH 8.0, 200 mM NaCl, 2 mM DTT, 50 mM maltose). Eluate was pooled and diluted 5× with Buffer D2 loaded onto a 5 ml HiTrap Heparin HP column (Cytiva) equilibrated in Buffer D2 + 75 mM NaCl. The column was washed with 5 CV Buffer D2 + 125 mM NaCl and eluted with a 20 CV 12.5-50% Buffer E2 gradient. The MBP tag was cleaved from full-length proteins (MEX3A-D, YTHDF, and mutants thereof) by overnight incubation with 200 µl 20 µM GST-3C protease with 2 ml amylose Sepharose. Flowthrough was used for further purification. For truncations of MEX3A-D, the MBP tag was not cleaved. Protein was pooled, concentrated, and loaded onto an S200 26/60 column (Cytiva), equilibrated in Buffer F2. Peak fractions were pooled, concentrated to ∼20 µM, flash frozen, and stored at -80°C.

Pellets expressing His_6_-SUMO tagged proteins (ZFP36 and MEX3C(418-600)) were resuspended in 2-3× Buffer A4 (50 mM HEPES pH 7.0, 500 mM NaCl, 2 mM DTT, 0.5 mM PMSF, 1× EDTA-free cOmplete protease inhibitor (Roche), 20 µg/ml DNase I) by volume, lysed, and centrifuged as above. Lysate was incubated with 6 ml equilibrated Ni^++^-NTA agarose resin (Qiagen) for 2 h, 4°C. Resin was washed with 50 ml Buffer A4, 50 ml Buffer B4 (50 mM HEPES pH 7.0, 1 M NaCl, 2 mM DTT), 50 ml Buffer C4 (50 mM HEPES pH 7.0, 200 mM NaCl, 10 mM imidazole, 2 mM DTT), and protein was eluted in 6 steps with 30 ml Buffer D4 (50 mM HEPES pH 7.0, 200 mM NaCl, 200 mM imidazole, 2 mM DTT). Eluate was diluted 8× in buffer E4 (25 mM HEPES pH 7.0, 1 mM DTT) and loaded onto a 5 ml S HP column (Cytiva). This was washed with buffer E4 + 75 mM NaCl and eluted with a 20 CV 7.5-50% Buffer F4 gradient (Buffer E4 + 1 M NaCl). Peak fractions were cleaved on 2 ml Ni^++^-NTA agarose beads with 200 µl 20 µM SUMO protease overnight, 4°C. Flowthrough was loaded on a 5 ml S HP column (Cytiva) and concentrated with a step elution to 50% Buffer F4. Peak fractions were loaded onto a S75 26/60 column, pre-equilibrated in Buffer G4 (20 mM HEPES pH 7.0, 200 mM NaCl, 1 mM TCEP). Fractions were pooled, concentrated to ∼20 µM, flash frozen, and stored at -80°C.

His-SUMO-tagged MEX3C(418-600) was purified as above for ZFP36, except flowthrough from SUMO protease cleavage was loaded onto a 5 ml Q HP column. Peak fractions were concentrated and loaded on a S200 16/60 column (Cytiva), pre-equilibrated in Buffer G. For pull-down studies, MEX3C(418-600)-lipoyl was pooled, concentrated, and flash frozen. For NMR studies, peak fractions containing MEX3C(418-600) from the S200 16/60 column were pooled and incubated overnight with 200 µl 20 µM GST-3C protease to cleave the C-terminal lipoyl tag, during dialysis against Buffer G4 + 0.02% azide. In addition to the amino acid sequence for MEX3C(418-600), the NMR construct contains a short linker (GS) and the 3C protease cleavage scar (LEVLFQ). The sample was concentrated and loaded onto an S75 10/300 column (Cytiva), peak fractions were pooled, concentrated, and used immediately for NMR studies.

Pellets expressing C-terminally TEV-SII-tagged PABPC1 were resuspended in 2-3× Buffer A5 (100 mM HEPES pH 8.0, 500 mM NaCl, 2 mM DTT, 20 µg/ml DNase I, 10 µg/ml RNase A, cOmplete protease inhibitors, 1 mM PMSF, 2 ml Biolock) by volume, lysed, and centrifuged as above. Lysate was incubated with 4 ml equilibrated StrepTactin resin (IBA) for 6 h, 4 °C to liberate PABPC1 from bound RNA. Resin was washed with 100 ml Buffer A5 and 100 ml Buffer B5 (50 mM HEPES pH 8.0, 1 M NaCl, 1 mM DTT), and protein eluted with 30 ml Buffer C5 (50 mM HEPES pH 8.0, 300 mM NaCl, 1 mM DTT, 20 mM DSB). Eluate was diluted 5× in Buffer D5 (25 mM HEPES pH 8.0, 1 mM DTT) and loaded onto a 5 ml HiTrap S HP column, This was washed with buffer D5 + 100 mM NaCl and eluted with a 25 CV 10-100% Buffer E5 gradient (D5 + 1 M NaCl). Peak fractions were digested with 200 µl 20 µM TEV protease overnight at 4 °C. This was diluted 8× in Buffer D and loaded onto a 5 ml HiTrap Heparin HP column. The column was washed with 10% Buffer E5, and protein was eluted with a 25 CV 10-100% Buffer E5 gradient. Fractions were concentrated and loaded onto a Superdex75 26/60 column (Cytiva), pre-equilibrated in Buffer F5 (20 mM HEPES pH 8.0, 200 mM NaCl, 1 mM TCEP). Peak fractions were pooled, concentrated to ∼50 µM, flash frozen, and stored at -80 °C.

### SDS PAGE Analysis

Protein samples were mixed with 4× LDS buffer (ThermoFisher) and denatured at 95°C, 5 min. All SDS-PAGE analysis was carried out using XCell SureLock Mini-Cell Systems. MOPS and MES running buffers were used for large and small proteins respectively. Gels were run at 200 V for 45-80 min depending on the required resolution. Gels were visualized using white light illumination with a GelDoc XR+ system (BioRad).

### *In vitro* pull-down assay

For pull-down assays between SII-tagged PAN2–PAN3 constructs and RNA-binding proteins, 50 µl 2 µM PAN2–PAN3-SII was immobilized on 15 µl StrepTactin (IBA) pre-incubated in pull-down buffer (PB, 10 mM HEPES pH 8.0, 200 mM NaCl, 1 mM TCEP) for 1 h, 4°C. The resin was washed 2× in 100 µl PB and incubated with 50 µl 4 µM prey protein for 2 h, 4 °C. 5 µl was removed as an input control (INPUT). The resin was resuspended and washed in 3× 100 µl PB and protein was eluted with 20 µl PB + 20 mM biotin. 15 µl of the eluate was removed (ELUATE). The input and eluate fractions were analyzed by SDS PAGE.

For pull-down assays between MBP-tagged MEX3C truncations and SII-tagged PAN2–PAN3, 50 µl 2 µM MBP-tagged MEX3C truncation was immobilized on 15 µl amylose sepharose resin (NEB) pre-incubated in PB for 1 h, 4°C. The resin was washed 2× in 100 µl PB and incubated with 50 µl 5 µM PAN2–PAN3 for 1 h, 4 °C. 5 µl was removed as an input control (INPUT). The resin was resuspended and washed in 3× 100 µl PB and protein was eluted with 20 µl PB + 50 mM maltose. 10 µl of the eluate was removed (ELUATE). The input and eluate fractions were analyzed by SDS PAGE.

For pull-down assays between SII-tagged PAN2–PAN3 truncations and MBP-MEX3C(418-600)-lipoyl, 50 µl 2 µM SII-tagged PAN2–PAN3 truncations was immobilized on 15 µl StrepTactin resin (IBA) pre-incubated in PB for 1 h, 4°C. The resin was washed 2× in 100 µl PB and incubated with 50 µl 5 µM MBP-MEX3C truncations for 2 h, 4 °C. 5 µl was removed as an input control (INPUT). The resin was resuspended and washed in 3× 100 µl PB and protein was eluted with 20 µl PB + 20 mM biotin. 5 µl of the eluate was removed (ELUATE). The input and eluate fractions were analyzed by SDS PAGE.

### *In vitro* deadenylation assay

*In vitro* deadenylation assays were carried out according to previous studies (Stowell et al. 2016; Webster et al. 2017) with slight modifications. In brief, RNA substrates containing an upstream region (optionally with a cognate RBP binding site) were designed to contain a 5′ fluorophore (6-FAM or Alexa647) and a 3′ 30-adenosine tail (60-adenosine tail for PABPC1 assays). 50 µl assays were performed at 37°C in Protein LoBind tubes (Eppendorf) in deadenylation buffer (20 mM PIPES pH 6.8, 10 mM KCl, 2 mM Mg(OAc)_2_, 0.1 mM TCEP). Deadenylase and RBPs (or no protein/tag alone for negative control) were diluted to 10× the indicated concentrations in 10 mM HEPES pH 8.0, 150 mM NaCl, 0.5 mM Mg(OAc)_2_, 0.5 mM TCEP. RBPs were incubated with 200 nM RNA for 1 h prior to the reaction. The reaction was topped up to 45 µl with DEPC H_2_O. Deadenylation reactions were initiated by addition of 5 µl deadenylase. 4 µl aliquots were removed at indicated time points and mixed with 4 µl formamide loading dye (96% formamide, 10 mM EDTA, 0.3% bromophenol blue, 1% SDS). Samples were resolved on a 20% urea denaturing polyacrylamide gel, run at 300 V for 60-75 min in 1× TBE. Fluorescently-labelled RNA was imaged on a Typhoon 5 Biomolecular Imaging System (excitation: 488 nm; emission filter: 520 nm) at 50 µm pixel size. Raw images were analyzed in Adobe Photoshop to linearly increase contrast.

### Two-color deadenylation assays

Two-color deadenylation assays were carried out as above with slight modifications. 100 nM 5′ 6-FAM-labelled or 5′ Alexa647-labelled RNAs with 30-adenosine tails were added to the assay, and 100 nM RBP was added instead of saturating concentrations. Deadenylation assays and denaturing gel analysis were carried out as above. Imaging was carried out on two channels (excitation: 488 nm; emission filter: 520 nm for 6-FAM; excitation: 635 nm; emission filter: 670 for Alexa647). Images were superimposed in Adobe Photoshop and false-colored for analysis.

### Electrophoretic mobility shift assays (EMSA)

EMSAs were carried out to determine the minimum RBP concentration required to saturate RNA. 10 µl binding reactions were prepared by adding protein at the indicated concentration to 200 nM 5′ 6-FAM-labelled RNA under deadenylation assay conditions (see above). The sample was incubated with 10× loading dye (20% glycerol, 0.25% Orange G) for 1 h to allow binding equilibrium to be reached. Samples were loaded onto 10% native TBE polyacrylamide gels and resolved by electrophoresis at 100 V, 1 h. Gels were scanned using a Typhoon 5 Biomolecular Imaging System (Cytiva) as above.

### NMR Spectroscopy

Experiments were performed on an in-house Bruker 700-MHz Avance II+ spectrometer equipped with a triple-resonance TCI CryoProbe. All NMR data were collected at 700 MHz in 10 mM HEPES pH 7.0, 200 mM NaCl, 1 mM TCEP, 0.02% NaN_3_, 5% D_2_O at 5°C.

Backbone resonance assignments of 100 µM MEX3C(418-600) were carried out using standard triple resonance experiments from the Bruker pulse sequence library, acquired with non-uniform sampling (15-30%) including HNCO, HN(CA)CO, HNCACB, CBCA(CO)NH, and HN(COCA)NNH. Resonances from proline residues were assigned using ^1^H start versions of ^13^C-detect CON, H(CA)CON, and H(CA)NCO experiments. Topspin 3.6.0 (Bruker) was used for data processing and involved compressed sensing (CS) reconstruction with qMDD (Mayzel et al., 2014) for sparse data sets. Backbone assignments were based on correlations of HN, N, Cα, Cβ and C’ chemical shifts in NMRFAM-Sparky 1.47 (Lee et al. 2015) supported with automatic chemical shift assignments in MARS (Jung & Zweckstetter, 2004) and curated manually using in-house scripts.

BEST (Band-selective Excitation Short Transient)-TROSY experiments were employed to enhance the sensitivity of titration experiments using 45 µM MEX3C(418-600). PAN3^PKC^ was added at either 1:20 or 1:10 (2.25 µM or 4.5 µM final concentrations). Weighted chemical shift perturbations of ^15^N-^1^H resonances were calculated using the equation (Ayed et al. 2001):

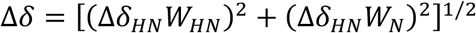

with weight factors determined from the average variances of chemical shifts in the BMRB chemical shift database (Mulder et al. 1999), where W_HN_ = 1 and W_N_ = 0.16.

### BioID Cell Line Preparation

Cell lines were generated in HEK293 Flp-In T-REx 293 (Invitrogen), grown at 37°C in high-glucose DMEM, supplemented with 5% Fetal Bovine Serum, 5% Cosmic calf serum and 100 U/ml Pen/Strep (growth media). Parental cell lines were routinely monitored for mycoplasma contamination.

To generate stable cell lines, Flp-In T-REx cells were transfected using jetPRIME transfection reagent (Polyplus Cat# CA89129-924). Cells were seeded at 250,000 cells/well in a 6-well plate in 2 ml growth media. The next day (day 1), cells were transfected with 200 ng pcDNA5-FLAG-BirA* or pcDNA5-FLAG-miniTurbo bait construct and 1 μg pOG44 in 200 μl of jetPRIME buffer mixed with 3 μl of jet Prime reagent , per manufacturers’ protocol. On day 2, transfected cells were passaged to 10 cm plates. On day 3, transfected cells were selected by adding 200 μg/ml hygromycin to the growth media. This selection media was changed every 2–3 days until clear visible colonies were present, at which point cells were scaled up to 150 mm plates.

Protein expression was induced using 1 μg/ml tetracycline when cells were at 80% confluence, and growth media was supplemented with 50 μM biotin for protein labeling (24 hours for BirA*, 20 min for miniTurbo). Cells were harvested by decanting growth media, washing with 5 ml PBS per 150 mm plate, and scraping in 1 ml ice-cold PBS. Cells from 1 × 150 mm plate were pelleted at 233 x g for 5 min, PBS was aspirated, and pellets were frozen on dry ice. Cell pellets were stored at −80°C until processing.

### BioID sample preparation

Cell pellets from two independent 150 mm plates were prepared as biological replicates. Samples were lysed using modRIPA-Lysis buffer at a 1 in 10 (mg:μl) ratio (50 mM Tris-HCl pH 7.5, 150 mM NaCl, 1% NP40, 1.5 mM MgCl_2_, 1 mM EGTA, 0.1% SDS, protease inhibitor cocktail (Sigma-Aldrich P8340, 1:500) and 0.4% sodium deoxycholate). Lysate was sonicated (three 5-second bursts with 2 seconds rest) on ice at 30% amplitude. Samples were digested with 250 U benzonase and RNaseA (ThermoFisher EN0531, 1 µg/ml) to reduce nucleic acid-mediated interactions (Sigma-Aldrich, E8263, 250U) for 10 min at 4°C with end-over-end rotation. SDS concentration was increased to 0.4%, followed by end-over-end rotation for 10 min at 4°C to further solubilize lysates, and clarified for 20 min at 20,817 × g at 4°C.

Streptavidin-sepharose beads (GE Cat# 17-5113-01) were washed 3 times with 1 ml RIPA buffer (50 mM Tris-HCl pH 7.5, 150 mM NaCl, 1 mM EDTA, 1% NP40, 0.1% SDS, 0.4% sodium deoxycholate). Beads were pelleted at 400 × g, 1 minute between washes. Clarified supernatants were incubated with 20 µl bed volume of washed beads in a fresh tube and rotated end-over-end for 3 hours at 4°C. Beads were pelleted (400 × g, 1 min), the supernatant was removed, and the beads were transferred to a new tube in 1 ml 2% SDS buffer (2% SDS, 50 mM Tris-HCl pH 7.5). The beads were washed 2× with 1 ml RIPA buffer, 1× in TENN lysis buffer (50 mM Tris-HCl pH 7.5, 150 mM NaCl, 1 mM EDTA, 0.1% NP-40), and 3× in 50 mM ammonium bicarbonate pH 8.0 (ABC). Beads were pelleted by centrifugation (400 × g, 1 min) and the supernatant was aspirated between wash steps. After the last wash, residual ABC was removed.

Beads were re-suspended in 60 μl of ABC with 1 μg trypsin and incubated at 37°C overnight with rotation. The next day, an additional 0.5 μg of trypsin was added to each sample (in 15 μl ABC), and the samples were incubated for an additional 3 hrs at 37°C with rotation. Beads were pelleted (400 × g, 2 min), and the supernatant was transferred to a fresh 1.5 ml tube. The beads were then rinsed 2× with 30 µl HPLC grade H_2_O (pelleting beads at 400 × g, 2 min in between); these rinses were combined with the original supernatant. The pooled supernatant was centrifuged at 16,100 × g, 10 min to remove beads and transferred to a new tube. 50% formic acid was added to pooled supernatant to a final concentration of 2% and dried using vacuum centrifugation. Dried peptides were stored at -80°C until mass spectrometry analysis.

### Mass spectrometry acquisition using TripleTOF mass spectrometers

BioID samples were subjected to mass spectrometry in two biological replicates. An eighth of each sample was loaded at 800 nl/min onto a 15 cm 100 μm ID emitter tip packed in-house with 3.5 μm Reprosil C18 (Dr. Maisch GmbH). Peptides were eluted from the column at 400 nl/min over a 90-minute gradient generated by a 425 NanoLC (Eksigent) and analyzed on a TripleTOF™ 6600 instrument (AB SCIEX). The gradient started at 2% acetonitrile with 0.1% formic acid and increased to 35% acetonitrile over 90 min followed by 15 min at 80% acetonitrile, and then 15 min at 2% acetonitrile. After each sample, the column was flushed twice for one hour at 1500 nl/min using an alternating five-minute sawtooth gradient from 35% acetonitrile to 80% acetonitrile to minimize carryover. Column and instrument performance was verified after each sample by analyzing a 30 fmol bovine serum albumin (BSA) tryptic peptide digest with a 60 fmol α-casein tryptic digest in a 30-minute gradient. Mass calibration for the mass spectrometer was performed on BSA reference ions between samples. Acquisition of ions was performed in data-dependent mode and consisted of one 250 ms MS1 TOF survey scan from 400-1250 Da followed by twenty 100 ms MS2 candidate ion scans from 100–2000 Da in high sensitivity mode. Ions with a charge of 2+ to 4+ exceeding the threshold of 200 counts per second were selected for fragmentation. Former precursors were excluded for 10s after one occurrence.

### Mass spectrometry peptide analysis and SAINT filtering

Data from the mass spectrometer were analyzed using the ProHits laboratory information management system (LIMS) platform (Liu et al. 2010). Wiff files were converted to MGF (WIFF2MGF) and to mzML format with ProteoWizard (3.0.4468) and AB SCIEX MS Data Converter (V1.3 beta). MzML files were searched using Mascot (v2.3.02) and Comet (2014.02 rev.2) (Eng et al. 2013). Search results were concatenated and analyzed with the Trans-Proteomic Pipeline (TPP) via the iProphet pipeline. Spectra were searched against 72,515 proteins consisting of the NCBI human RefSeq database (version 55; 27 March 2020; forward and reverse sequences), and ‘common contaminants’ from the Max Planck Institute (https://lotus1.gwdg.de/mpg/mmbc/maxquant_input.nsf/7994124a4298328fc125748d0048fee2/$FILE/contaminants.fasta), Global Proteome Machine (http://www.thegpm.org/crap/index.html), and sequences from common fusion proteins and epitope tags. Database parameters were adjusted to search for tryptic cleavages, allowing up to two missed cleavage sites per peptide, MS1 mass tolerance of 40 ppm with charges of 2+ to 4+, and an MS2 mass tolerance of ± 0.15 amu. Asparagine/glutamine deamidation and methionine oxidation were selected as variable modifications. A minimum iProphet probability of 0.95 was required for protein identification. Proteins detected with a minimal number of two unique peptides were used for protein interaction scoring.

Significance Analysis of INTeractome (SAINTexpress version 3.6.1) was applied to calculate the probability of potential interactions from the background (Teo et al. 2014). Biological replicates for N- and C-terminal-tagged baits were analyzed against a corresponding set of control samples (untransfected cells or cells expressing BirA*- FLAG/miniTurbo-FLAG, or BirA*-FLAG-GFP/miniTurbo-FLAG-GFP). Each biological replicate was analyzed independently against the control before averaging biological replicate results and calculating Bayesian False Discovery Rates (BFDR). High-confidence interactions were those with BFDR ≤ 0.01.

### Data Analysis and Visualization using ProHits-viz

Proximal protein-protein interactions scored with SAINTexpress were used to generate input files for ProHits-viz (Knight et al. 2017). Prey-centric identification of PAN2 and PAN3 proximal interactors were carried out using data from **Supplementary Table 1** from Youn and colleagues (Youn et al. 2018). Bait proteins which biotinylated PAN2 and PAN3 were filtered by removal of duplicates, BFDR ≤ 0.01, and 20× fold enrichment over the negative control (PAN2: n=26; PAN3: n=22). Bait proteins proximal to both PAN2 and PAN3 and satisfying the above criteria (n=20) were prioritized, annotated by known function, and visualized using a simplified network diagram.

Unless otherwise indicated, SAINTexpress output files for labeled PAN2, PAN3, or PAN3^PKC^ (**Table S2**) were filtered by BFDR ≤ 0.1. To generate area-proportional Venn diagrams (**Figure 5b, 6a**), proximal interactors were filtered, annotated by function, and resultant proximal interactors were compared. To generate bait-bait scatter plots (**Figure 5c, 6b, S8b-c**), control subtraction and data normalization were carried out from SAINTexpress outputs by default. Prey proteins (BFDR ≤ 0.01) were quantified by average spectral count in each condition. Previously known interactors, newly identified interactors from this study, and putative RBP interactors are colored and labeled accordingly. To generate dot plots, only prey proteins present across all three (**Figure S8d**) or four (**Figure 5a**) tag conditions (PAN3 and PAN3^PKC^) were considered. Default options for control subtraction and data normalization were selected. Dot color is scaled from light blue to black to the average spectral count (AvgSpec) across each tagged construct; dot area is scaled according to AvgSpec between prey proteins; outer ring color was determined by BFDR. Data were arranged manually where stated (spectral counts summed and divided by two) and annotated according to function.

### Bioinformatic Analysis

Sequence alignment was carried out using the T-Coffee suite (Notredame et al. 2000) and visualized using EsPript 3.0; alignments were colored based on percentage sequence similarity (Robert and Gouet 2014). Disorder prediction of proteins of interest was carried out using DISOPRED3 (Jones and Cozzetto 2015) and visualized using GraphPad Prism 9.3.0 (GraphPad Software). Proteins from Mass Spectrometry data were functionally annotated for RNA-binding using data from the Gene Ontology Project (Baltz et al. 2012; Castello et al. 2012; Gaudet et al. 2011). Expression levels of proteins of interest in HEK293 cells were obtained from PaxDB (Wang et al. 2015), and confirmed with raw data (Geiger et al. 2012).

### Diagrams and Schematics

All figures were created and arranged in Adobe Illustrator. Proportional Venn diagrams were made using DeepVenn and subsequently modified (Hulsen et al. 2008).

## SUPPLEMENTARY FIGURES

**Table S1. Identification of high-confidence putative interactors of the PAN2–PAN3 complex by prey-centric analysis.**

Raw data were obtained from Table S1, Sheet (B) from Youn et al., 2018. Bait proteins which were proximal to (A) PAN2 and (B) PAN3 were obtained. These were filtered by fold-enrichment ≥ 20 and BFDR ≤ 0.01 to obtain high-confidence proximal partners (C). These proximal partners were annotated by known function (color), and overlap with both PAN2 and PAN3 (red text).

Columns A-B: bait gene name and position of the BirA*-FLAG tag (N or C). Columns C-D: prey protein ID (C) and prey gene name (D) derived from NCBI-assigned accession ID and name for PAN2 (sheet A) and PAN3 (sheet B). Columns E-I: spectral counts for PAN2 or PAN3 (E, delimited by “|”); spectral sum (F); averaged spectral counts (G); number of replicates (H); and spectral counts for PAN2 or PAN3 prey across negative controls (I). Columns J-K: averaged across replicates (J) and maximal (K) of all individual SAINT probability scores. Column L: pull-down counts divided by the control counts plus a small factor to prevent division by 0. Column M: Bayesian False Discovery Rate (BFDR) for each putative interactor with PAN2 or PAN3. All columns are derived directly from the SAINTexpress output; bait proteins are sorted by FoldChange in descending order.

Data from Table S1 were used to generate **Figures 1a** and S1a.

**Table S2. High-confidence BioID proximity interactome of the human PAN2–PAN3 deadenylase complex**

SAINTexpress Task IDs 5658, 5659, 5674, and 5676. In brief, PAN2, PAN3 or PAN3_PKC (PAN3^PKC^) constructs were N- (“BirA-”; “mT-”) or C- (“-BirA”; “-mT”) terminally tagged with BirA*-FLAG (A) or miniTurbo-FLAG (B). High-confidence proximal interactor prey proteins were obtained by averaging the N- and C-terminal BirA* (C) and miniTurbo (D) spectral counts respectively.

Column A: bait gene name or construct (PAN2, PAN3, or PAN3_PKC) and position of the BirA*-FLAG or miniTurbo-FLAG tags (N- or C-terminal), where applicable. Columns B-C: prey protein ID (B) and prey gene name (C) derived from NCBI-assigned accession ID and name. Columns D-H: spectral counts for prey proteins (D, delimited by “|”); spectral sum (E); averaged spectral counts (F); number of replicates (G); and spectral counts for PAN2 or PAN3 prey across negative controls (H). For Sheets C-D, the averaged spectral counts (column F) represent the average across N- and C-terminal tags. Columns I-J: averaged across replicates (I) and maximal (J) of all individual SAINT probability scores. Columns K-L: averaged across replicates (K) and maximal (L) of all individual topology-assisted SAINT probability scores. Column M: SaintScore: larger of AvgP and TopoAvgP. In this analysis, for all prey proteins, AvgP = TopoAvgP. Column N: logOddsScore: log likelihood ratio of the observed data over the expected value. Column O: pull-down counts divided by the control counts plus a small factor to prevent division by 0. Column P: Bayesian False Discovery Rate (BFDR) for each putative interactor with PAN2 or PAN3. All columns are derived directly from the SAINTexpress output; prey proteins are sorted by their accession ID.

Data from **Table S2** were used to generate **Table S3**. Proximity interactome data for PAN3 and PAN3^PKC^ were filtered for Bayesian False Discovery Rate (BFDR) < 0.1 and used to generate **Figures 5a** and **S8d**.

**Table S3. Annotated analysis of BioID proximity interactome of the human PAN2–PAN3 deadenylase complex and comparison with human CCR4–NOT complex**

In all sheets: column A: bait gene name or construct (PAN2, PAN3, or PAN3_PKC) and position of the BirA*-FLAG or miniTurbo-FLAG tags (N- or C-terminal), where applicable. Columns B-C: prey protein ID (B) and prey gene name (C) derived from NCBI-assigned accession ID and name.

Columns D-H: spectral counts for prey proteins (D, delimited by “|”); spectral sum (E); averaged spectral counts (F); number of replicates (G); and spectral counts for PAN2 or PAN3 prey across negative controls (H). For Sheets C-D, the averaged spectral counts (column F) represent the average across N- and C-terminal tags. Columns I-J: averaged across replicates (I) and maximal (J) of all individual SAINT probability scores. Columns K-L: averaged across replicates (K) and maximal (L) of all individual topology-assisted SAINT probability scores. Column M: SaintScore: larger of AvgP and TopoAvgP. In this analysis, for all prey proteins, AvgP = TopoAvgP. Column N: logOddsScore: log likelihood ratio of the observed data over the expected value. Column O: pull-down counts divided by the control counts plus a small factor to prevent division by 0. Column P: Bayesian False Discovery Rate (BFDR) for each putative interactor with PAN2 or PAN3. All columns are derived directly from the SAINTexpress output; prey proteins are sorted by their accession ID.

Sheets which are annotated: column R: if the prey protein is annotated as an RBP; column S: if the annotated RBP has known RNA sequence specificity/preference; column T: localization of the prey protein; column U: annotated function of the prey protein. UNCHAR: uncharacterized RNA-binding specificity or function. Annotations were only carried out for highlighted prey proteins.

**(A-B) Comparison of proximal prey proteins of PAN2 and PAN3.** N- and C-terminally averaged BirA*- (A) or miniTurbo- (B) tagged PAN2 and PAN3 were compared to validate newly identified interactors of the PAN2–PAN3 complex *in cellulo*. Datasets (PAN2-BirA: n=808; PAN3-BirA: n=612; PAN2-mT: n=341; PAN3-mT: n=311) were filtered by Bayesian False Discovery Rate (BFDR) < 0.1 (PAN2-BirA: n=70; PAN3-BirA: n=124; PAN2-mT: n=75; PAN3-mT: n=124). Prey proteins which were proximal to PAN2 and PAN3 were highlighted and annotated. These sheets were used to generate **Figure S8b**.

**(C-E) Comparison of proximal prey proteins of BirA* and miniTurbo tagged PAN2, PAN3, or PAN3^PKC^.** N- and C-terminally averaged BirA*- or miniTurbo-tagged (C) PAN2; (D) PAN3; or (E) PAN3^PKC^ were compared to obtain high-confidence proximity interactors of each construct. Datasets (PAN2-BirA: n=808; PAN2-mT: n=341; PAN3-BirA: n=612; PAN3-mT: n=311; PAN3^PKC^-BirA: n=903; PAN3^PKC^-mT: n=437) were filtered by Bayesian False Discovery Rate (BFDR) < 0.1 (PAN2-BirA: n=70; PAN2-mT: n=75; PAN3-BirA: n=124; PAN3-mT: n=124; PAN3^PKC^-BirA: n=58; PAN3^PKC^-mT: n=155). Prey proteins which were proximal to both BirA*-tagged and miniTurbo-tagged PAN2, PAN3, or PAN3^PKC^ constructs were highlighted and annotated in each sheet. These sheets were used to generate **Figure S8c**.

**(F) Comparison of high-confidence proximal interactors of PAN3 and PAN3^PKC^.** N- and C-terminally averaged BirA*- or miniTurbo-tagged PAN3 and PAN3^PKC^ were compared to identify shared prioritized proximity interactors of each construct. Filtered interactomes from **Table S3D-E** were obtained and putative RBP prey interactors (green), putative non-RBP prey interactors (red), and exclusive PAN3-proximal (yellow) or exclusive PAN3^PKC^-proximal (orange) prey interactors were identified and highlighted. This sheet was used to generate **Figure 5b-c**.

**(G) Comparison of high-confidence proximal RBP interactors of PAN3 and CNOT9.** Prioritized high-confidence proximal RBP interactors of PAN3 (n=46) were compared to those of CNOT9, obtained from our previous proximal interactome dataset (Youn et al., 2018). The BirA*-tagged CNOT9 interactome was N- and C-terminally averaged (n=1787) was filtered by Bayesian False Discovery Rate (BFDR) < 0.1 (n=236). Only prey proteins common to N- and C-terminal tags were considered (n=99) and were subsequently selected for known RNA-binding or RNA-processing function (n=49). This curated list of prioritized RBP interactors of the CCR4–NOT complex was compared to that of PAN3. Prey proteins common to both PAN3 and CNOT9 were highlighted. This sheet was used to generate **Figure 6a-b**.

---

**Figure S1.**
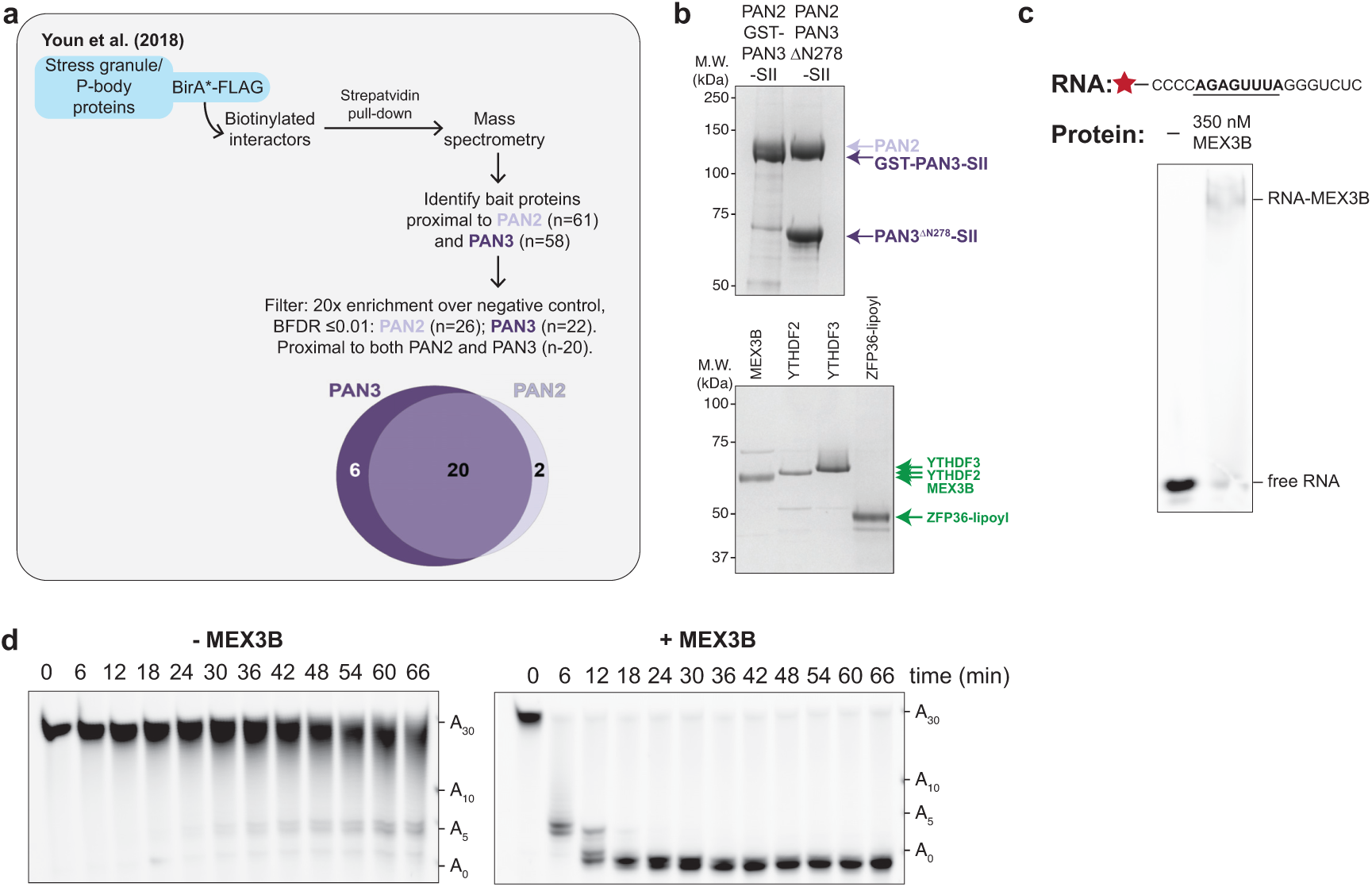
MEX3B interacts with and promotes targeted deadenylation by PAN2–PAN3. (a) **Schematic of prey-centric analysis of BioID data from Youn et al. (2018).** In that study, components of P-bodies and stress granules were tagged with BirA*-FLAG. Pull-down and mass spectrometry identified putative interactors, including PAN2 and PAN3 (**Table S1)**. BirA*-tagged bait proteins proximal to PAN2 (purple, n=61) and PAN3 (lilac, n=58) were filtered, resulting in a list of unique high-confidence interactors of PAN2 (n=26) and PAN3 (n=22). Bait proteins satisfying these criteria and proximal to both proteins (n=20) were further analyzed here. **(b) Purification of *H. sapiens* PAN2–PAN3 and RNA-binding proteins (RBPs).** PAN2 (lilac)–PAN3 (purple; and truncations thereof) and RBPs (green) were expressed (in *Sf*9 cells and *E. coli*, respectively), purified and analyzed by SDS-PAGE. **(c) Interaction of MEX3B with target RNAs *in vitro*.** Electrophoretic mobility shift assay (EMSA) of MEX3B with a synthetic target RNA, resolved by native PAGE. **(d) MEX3B accelerates deadenylation by full-length PAN2–PAN3 *in vitro*.** Synthetic RNA substrate containing a 30-nt poly(A) tail downstream of non-poly(A) ribonucleotides containing the MEX3 Recognition Element (MRE) was incubated at a saturating RBP-RNA ratio prior to adding 40 nM full-length wild-type PAN2–PAN3. Reactions were analyzed by denaturing PAGE. Substrate sizes with various poly(A) tail lengths are indicated.

**Figure S2.**
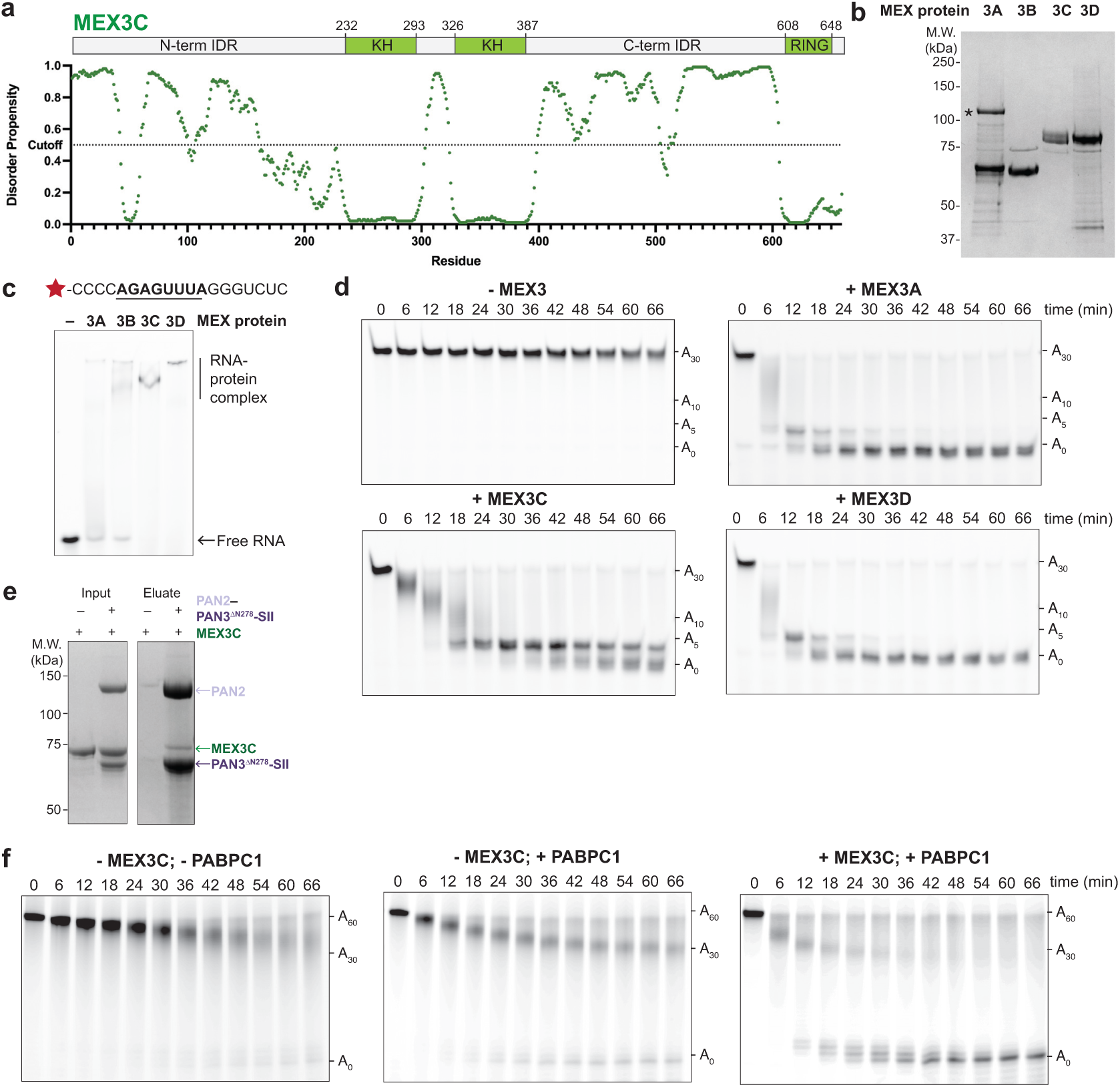
MEX3 paralogs interact with PAN2–PAN3 and promote deadenylation. (a) **Domain schematic of MEX3C.** MEX3C comprises an N-terminal intrinsically disordered region (IDR), two tandem KH domains which bind a specific RNA sequence, a C-terminal IDR, and a RING domain at the extreme C-terminus. A plot of disorder propensity against amino acid number (DISOPRED3) demonstrates negative correlation between disorder propensity and known domains. The dotted line at 0.5 denotes an arbitrary cutoff for disorder propensity. **(b) Recombinant MEX3A-D.** The purified proteins were resolved on SDS-PAGE. The asterisk indicates a major impurity which was present following purification of MEX3A. Molecular weights are indicated. **(c) MEX3 paralogs bind the MRE RNA.** Electrophoretic mobility shift assays (EMSAs) were used to test 350 nM MEX3 paralog binding to fluorescently labelled RNAs containing the cognate MRE (bold, underlined). Samples were resolved by native PAGE. **(d) MEX3 paralogs promote deadenylation by PAN2–PAN3.** Deadenylation assays were carried out with RNA substrates containing 19 non-poly(A) ribonucleotides with the MEX3 Recognition Element (MRE; shown in (c)) followed by a 30-adenosine poly(A) tail. MEX3A, MEX3C, and MEX3D, were added to MRE RNA prior to adding 40 nM PAN2–PAN3^ΔN278^-SII. Reactions were analyzed by denaturing PAGE. RNA sizes with various poly(A) tail lengths are indicated. **(e) MEX3C directly interacts with PAN2–PAN3.** Pull-down assays were carried out using purified MEX3C, and PAN2–PAN3^ΔN278^-SII as bait immobilized on StrepTactin resin, or a negative control (no immobilized protein). Input: immobilized protein + MEX3C in solution; eluate: biotin elution. Resulting samples were resolved by SDS-PAGE, and molecular weights are indicated. **(f) MEX3 increases the deadenylation rate by PAN2–PAN3 on a PABPC1-bound poly(A) tail.** Deadenylation assays were carried out with 200 nM MRE RNA substrates and a 60-adenosine poly(A) tail, in the presence or absence of 350 nM poly(A)-binding protein PABPC1. In the right panel, MEX3C was added to saturate MRE RNA prior to adding 40 nM PAN2–PAN3^ΔN278^-SII. Reactions were analyzed by denaturing PAGE. RNA sizes with various poly(A) tail lengths are indicated. Fully deadenylated RNA appears at earlier time points when both PABPC1 and MEX3C are present.

**Figure S3.**
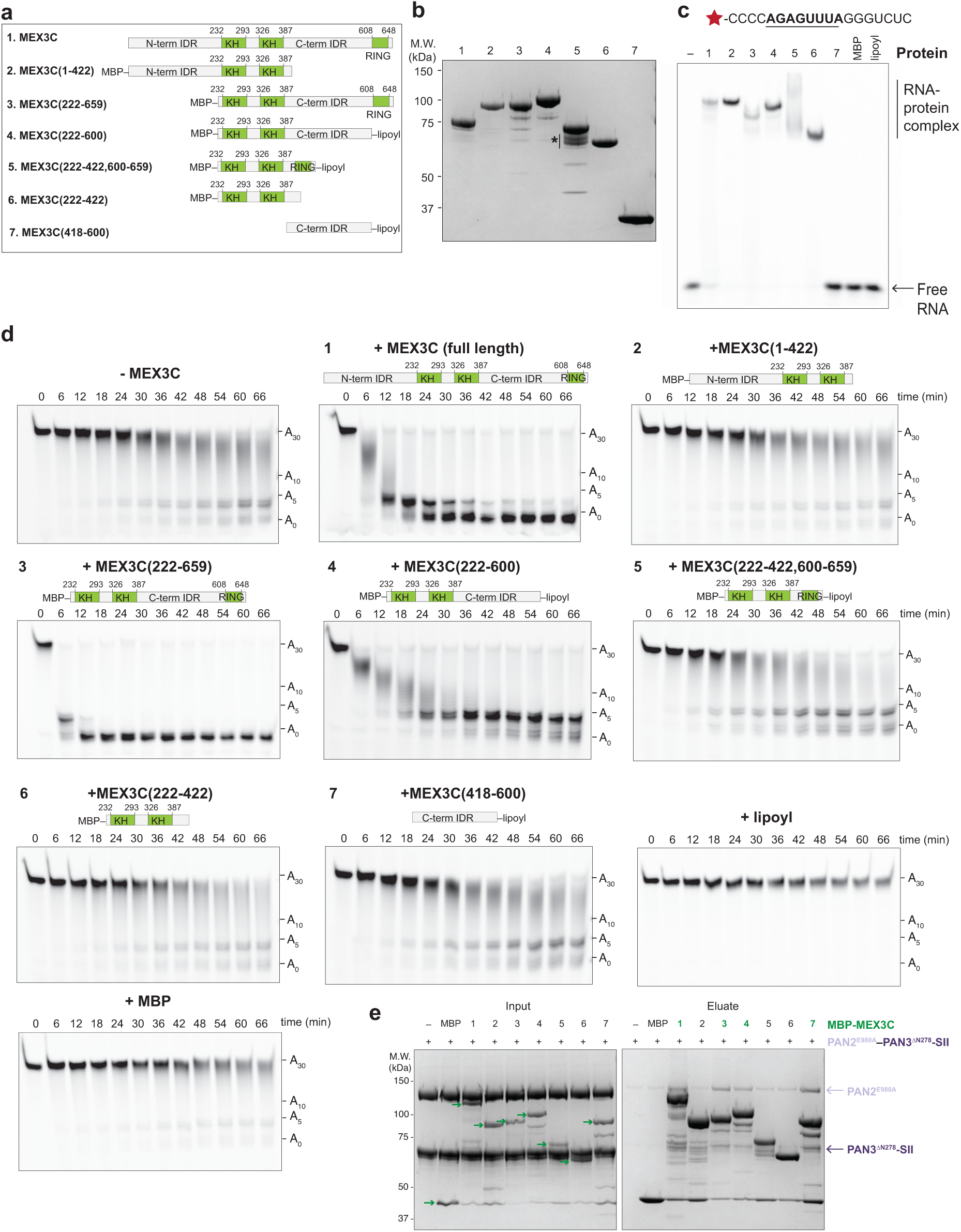
The C-terminal IDR of MEX3 interacts with and contributes to the acceleration of PAN2– PAN3 in targeted deadenylation. (a) **Diagram of MEX3C truncation constructs.** To determine which region of MEX3C is required for its ability to promote deadenylation, we designed constructs which contained the RNA-binding KH domains (residues 222-422) in tandem with other regions of the protein including: the N-terminal IDR (1-422), the entire C-terminal region (222-659), the C-terminal IDR (222-600), or the C-terminal RING domain (222-422,600-659). Some constructs included N-terminal MBP and/or C-terminal lipoyl tags as indicated. **(b) Truncation constructs of MEX3C.** Proteins were expressed in *E. coli*, purified and analyzed by SDS-PAGE. Constructs are numbered as in (a). The asterisk indicates impurities following purification, which are likely degradation products of the full-length protein. Molecular weights are indicated. **(c) MEX3C truncations containing the tandem KH domains bind RNA.** Electrophoretic mobility shift assays (EMSAs) were used to test binding of MEX3C truncations to 200 nM fluorescently-labelled RNAs containing the MRE (bold, underlined). Proteins are numbered according to panel (a). The IDR(418-600) alone (construct 7), MBP, and the lipoyl domain did not bind RNA. **(d) The C-terminal IDR of MEX3C contributes to acceleration of PAN2–PAN3 deadenylation.** Deadenylation assays were performed with the MRE RNA containing a 30-adenosine poly(A) tail and MEX3C variants or tags only (MBP, lipoyl) with 40 nM PAN2–PAN3^ΔN278^-SII. Reactions were analyzed by denaturing PAGE. RNA sizes with various poly(A) tail lengths are indicated. Affinity or solubility tags alone, the KH domains (222-422; construct 6) and the MEX3C C-terminal IDR alone (418-600; construct 7) do not substantially affect deadenylation. The full C-terminal region of MEX3C (construct 3) promotes deadenylation similarly to the full-length protein (construct 1) whereas the N-terminal IDR (construct 2) has a minimal effect. A construct containing the C-terminal IDR but lacking the RING finger domain (residues 222-600; construct 4) also stimulates deadenylation. **(e) The MEX3C C-terminal IDR directly interacts with PAN2–PAN3.** Pull-down assays were carried out using purified PAN2–PAN3^ΔN278^-SII as prey (lilac and purple arrows respectively), and N-terminal MBP-tagged MEX3C truncations as bait (green arrows), immobilized on amylose resin, or negative controls where no protein or MBP was immobilized. Input: immobilized protein + PAN2–PAN3^ΔN278^-SII in solution; eluate: maltose elution. Bait proteins are numbered according to the key in panel (a). Resulting samples were analyzed by SDS-PAGE, and molecular weights are indicated. Constructs which pull down PAN2–PAN3 above the negative control (1, 3, 4 and 7) contain the C-terminal IDR and are labelled in green. The MEX3C C-terminal IDR (residues 418-600; construct 7) directly interacts with PAN2–PAN3, but constructs lacking the C-terminal IDR have a negligible interaction. Overall, these data show that the C-terminal IDR of MEX3C directly interacts with PAN2-PAN3 and contributes to MEX3C-targeted poly(A) removal.

**Figure S4.**
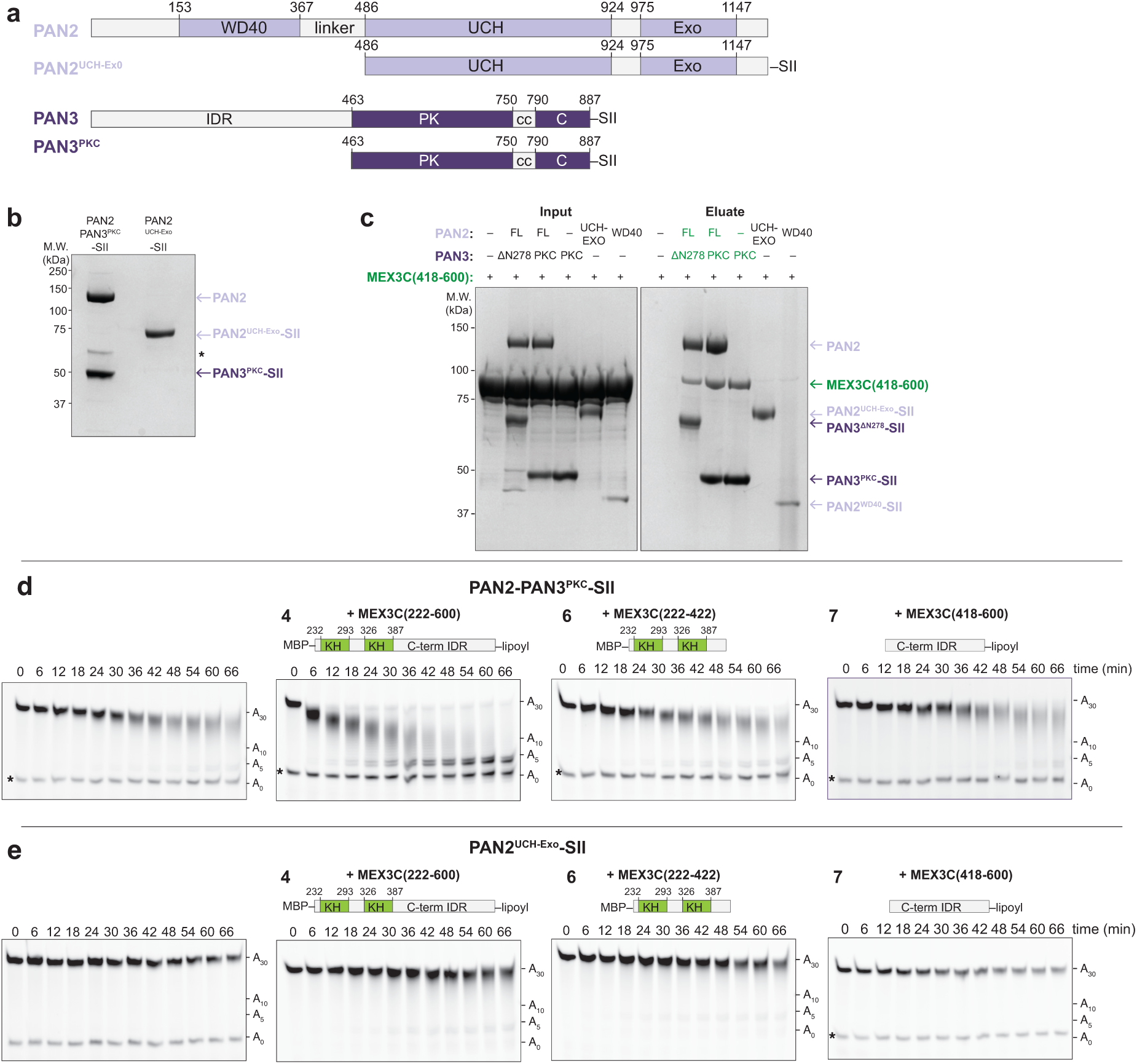
The C-terminal region of PAN3 is required for interaction with MEX3C and increased PAN2–PAN3 deadenylation activity. (a) **Domain schematics of PAN2 and PAN3.** PAN2 (lilac) contains an N-terminal WD40 domain, a linker which wraps around the PAN3 dimer, and a C-terminal catalytic unit consisting of structurally contiguous ubiquitin hydrolase and exonuclease domains (UCH-Exo, residues 516-1202) (Jonas et al. 2014; Schäfer et al. 2014; Wolf et al. 2014). PAN3 (purple) contains an N-terminal IDR, followed by contiguous pseudokinase (PK), coiled-coil (c), and C-terminal knob (C) domains (together PKC, residues 463-887), which mediate PAN3 dimerization (Christie et al. 2013; Wolf et al. 2014). Constructs used in deadenylation assays are shown. **(b) Purified PAN2–PAN3 truncation constructs.** PAN2–PAN3^PKC^-SII and PAN2^UCH-Exo^-SII were expressed in *Sf*9 insect cells, purified and resolved on SDS-PAGE to demonstrate their purity. The asterisk indicates an impurity which was present following purification of PAN2–PAN3^PKC^-SII. Molecular weights are indicated. **(c) The PAN3 PKC domains directly interact with the MEX3C C-terminal IDR.** SDS-PAGE of pull-down assays carried out using purified MBP- and lipoyl-tagged C-terminal IDR of MEX3C (418-600) and immobilized PAN2–PAN3 truncations containing an SII tag, or a negative control (no immobilized protein). Constructs of PAN2 alone (WD40 and UCH-Exo) did not interact with the MEX3C C-terminal IDR. Input: immobilized protein + MBP-MEX3C(418-600)-lipoyl in solution; eluate: biotin elution. **(d,e) The PAN3 PKC domains are required for MEX3C-mediated acceleration of PAN2–PAN3 deadenylation.** Deadenylation assays of an RNA substrate containing the MEX3 Recognition Element and a 30-adenosine poly(A) tail with no MEX3C or with the indicated MEX3C truncation constructs. The MEX3C constructs are numbered as in Figure S3a. Reactions with 40 nM PAN2–PAN3^PKC^-SII (d) or PAN2^UCH-Exo^ (e) were analyzed by denaturing PAGE. MEX3C KH domains or C-terminal IDR alone do not promote deadenylation. The sizes of the RNA substrate with various poly(A) tail lengths are indicated. Asterisks indicate an RNA degradation product that was present prior to the addition of PAN2–PAN3.

**Figure S5.**
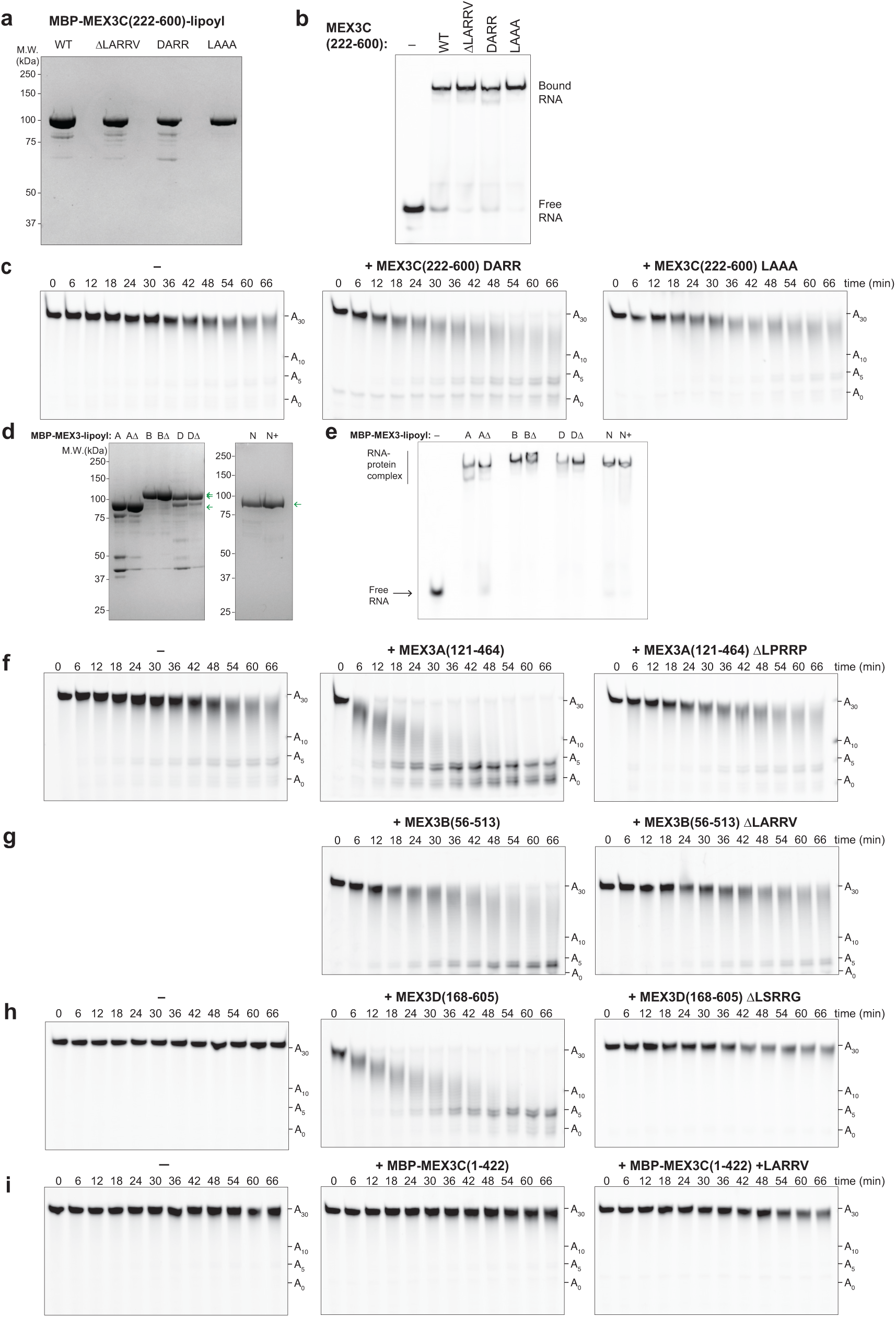
The LARR motif plays a role in PAN2–PAN3 targeted deadenylation. (a) **MEX3C LARR mutants.** LARR motif mutants of the MEX3C(222-600) construct, with N-terminal MBP and C-terminal lipoyl tags, were expressed in *E. coli*, purified and analyzed on SDS-PAGE. Mutants include deletion of LARRV (ΔLARRV), L556D (DARR) and R558A/R559A (LAAA). Molecular weights are indicated. **(b) MEX3C LARR mutants bind RNA.** Electrophoretic mobility shift assays (EMSAs) were used to assess binding of 350 nM MEX3C(222-600) LARR mutants to 200 nM fluorescently-labelled RNAs containing the MRE. Mutant proteins fully shifted the RNA similar to wild-type. **(c) Mutations in the LARR motif attenuate MEX3C acceleration of PAN2-PAN3 targeted deadenylation.** 350 nM DARR (middle) and LAAA (right) mutants of MBP-MEX3C(222-600)-lipoyl were added to deadenylation reactions containing 40 nM PAN2–PAN3^ΔN278^-SII. Reactions were analyzed by denaturing PAGE. The sizes of the substrate with various poly(A) tail lengths are indicated. Compare to Figure 2c. **(d) Human MEX3 paralogs.** All constructs were expressed in *E. coli*, purified and analyzed by SDS-PAGE. Constructs include the RNA-binding KH domains and the N- or C-terminal IDRs. A(Δ): MEX3A(121-464) (ΔLPRRP); B(Δ): MEX3B(56-513) (ΔLARRV); D(Δ): MEX3D(168-605) (ΔLSRRG); N(+): MEX3C(1-422) (+LARRV after residue 73). Green arrows indicate the full-length proteins, which contain N-terminal MBP and C-terminal lipoyl tags. **(e) LARR motif mutants bind MRE-containing RNA.** Electrophoretic mobility shift assays (EMSAs) were used to assess 350 nM MEX3 paralogs (wild-type and LARR deletions), and MEX3C(1-422) with an inserted LARRV motif binding to 200 nM MRE-containing RNA. **(f-h) The LARR motifs of MEX3 paralogs contributes to targeted deadenylation by PAN2–PAN3.** MEX3A (f), MEX3B (g), and MEX3D (h) constructs containing or lacking (Δ) the conserved LARR motif were incubated with MRE-containing RNAs with a 30-nt poly(A) tail at saturating concentrations and used in deadenylation assays with PAN2–PAN3^Δ278^-SII (f,g 60 nM; h, 20 nM). Reactions were analyzed by denaturing PAGE. RNA sizes with various poly(A) tail lengths are indicated. Each wild-type construct accelerated PAN2– PAN3 deadenylation but to different degrees. For all paralogs, deletion of the LARR motif impaired PAN2– PAN3 deadenylation, suggesting that this motif is generally important in MEX3 function. **(i) The LARR motif is not sufficient to promote PAN2–PAN3 activity.** LARRV was introduced into MEX3C(1-422) at a similar distance in sequence to the RNA-binding KH domains as in wild-type. The RBPs were incubated with MRE-containing RNAs with a 30-nt poly(A) tail at saturating concentrations and a deadenylation reaction with 20 nM PAN2–PAN3^Δ278^-SII was performed. Reactions were analyzed by denaturing PAGE. RNA sizes with various poly(A) tail lengths are indicated.

**Figure S6.**
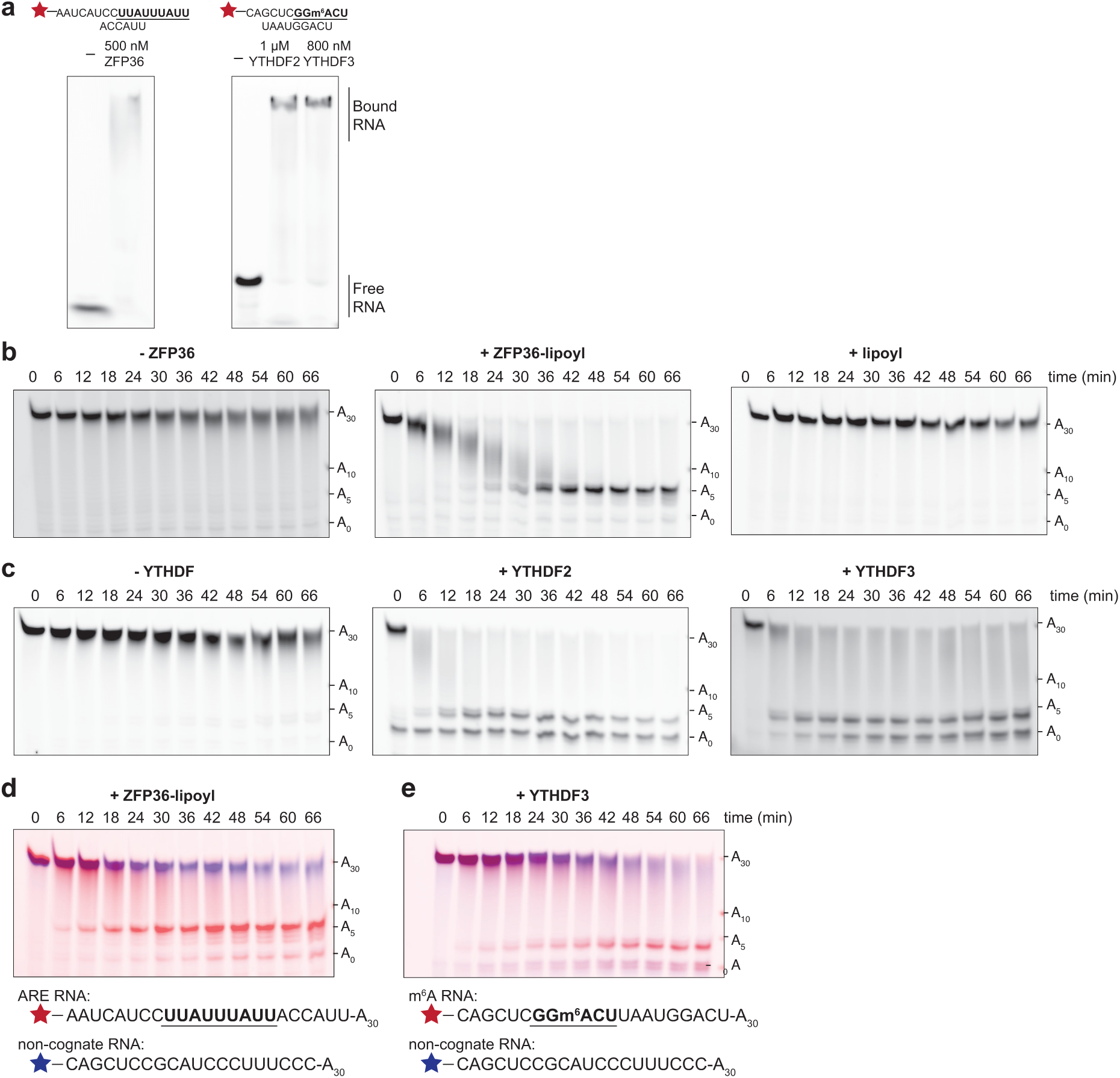
Additional RNA-binding proteins interact with and promote targeted deadenylation by PAN2– PAN3. (a) **Interaction of RBPs with model RNAs *in vitro*.** Electrophoretic mobility shift assays (EMSAs) of ZFP36, YTHDF2 and YTHDF3 with model RNAs, resolved by native PAGE. RNA sequences are indicated. **(b-c) ZFP36, YTHDF2 and YTHDF3 accelerate deadenylation by full-length PAN2–PAN3 *in vitro*.** Synthetic RNA substrates containing a 30-nt poly(A) tail downstream of non-poly(A) ribonucleotides containing the RBP motifs were incubated at a saturating RBP-RNA ratio prior to adding 40 nM full-length wild-type PAN2–PAN3. The lipoyl domain alone was included as a negative control. Reactions were analyzed by denaturing PAGE. Substrate sizes with various poly(A) tail lengths are indicated. **(d-e) ZFP36, YTHDF2 and YTHDF3 mediate selective targeted deadenylation *in vitro*.** Deadenylation of an equimolar mixture of 100 nM fluorescein-labelled target RNA (red) and 100 nM Alexa647-labelled non-cognate RNA (blue) containing a 30-nt poly(A) tail with (d) 20 nM PAN2–PAN3^ΔN278-SII^ or (e) 40 nM PAN2– PAN3^ΔN278-SII^. The RBP binding motifs are in bold and underlined. A 1:1 RBP:cognate RNA ratio was used to maximize specific (cognate) interactions over non-specific (non-cognate) interactions.

**Figure S7.**
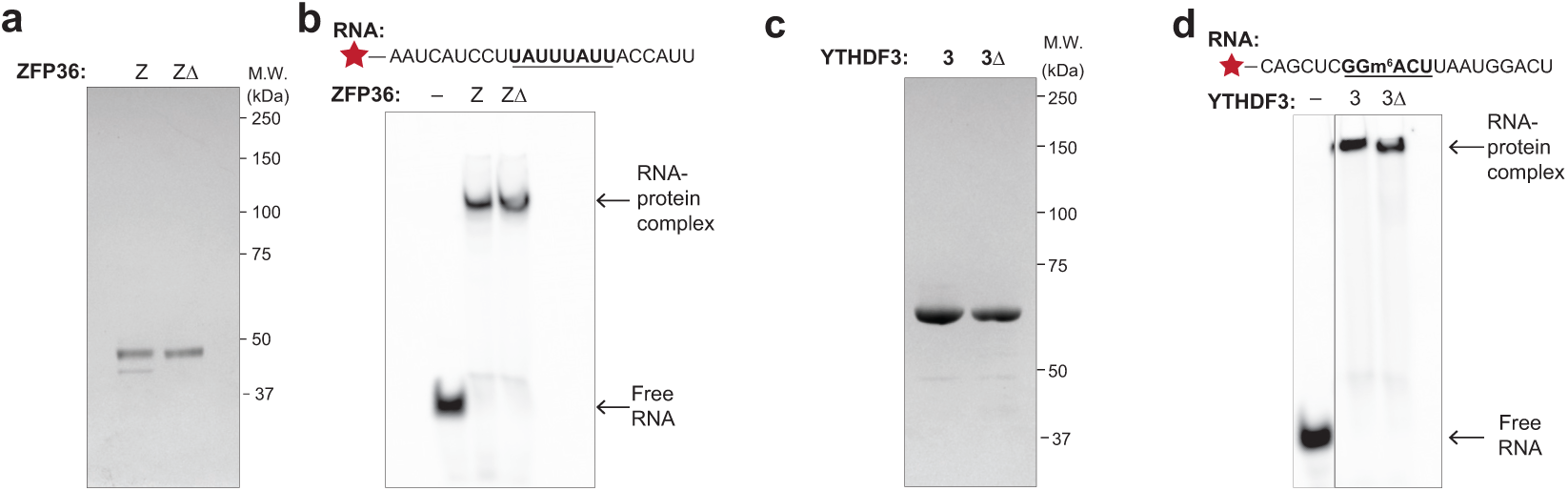
ZFP36 and YTHDF3 LARR-motif mutants. (a) **SDS PAGE of wild-type ZFP36 (Z) and a construct lacking LARRD (ZΔ).** The proteins were expressed in *E. coli*, purified and analyzed by SDS-PAGE. Molecular weights are indicated. **(b) Constructs of ZFP36 containing and lacking the LARRD motif bind RNA.** Fluorescently labelled RNAs containing the consensus ARE were incubated with 500 nM ZFP36 (Z), or 500 nM ZFP36 ΔLARRD (ZΔ). RNA binding was assessed by electrophoretic mobility shift assays (EMSAs). **(c) Purification of YTHDF3 mutants.** SDS PAGE analysis of YTHDF3 (3), and YTHDF3 lacking the IARK motif (3Δ), expressed in *E. coli*, purified and analyzed on SDS-PAGE. Molecular weights are indicated. **(d) YTHDF3 proteins containing and lacking the IARK motif bind RNA.** Fluorescently labelled RNAs containing the GGm^6^ACU YTHDF consensus RNA-binding motif were incubated with 800 nM YTHDF3 (3), or 800 nM YTHDF3 ΔIARK (3Δ) and RNA binding was assessed by electrophoretic mobility shift assays (EMSAs).

**Figure S8.**
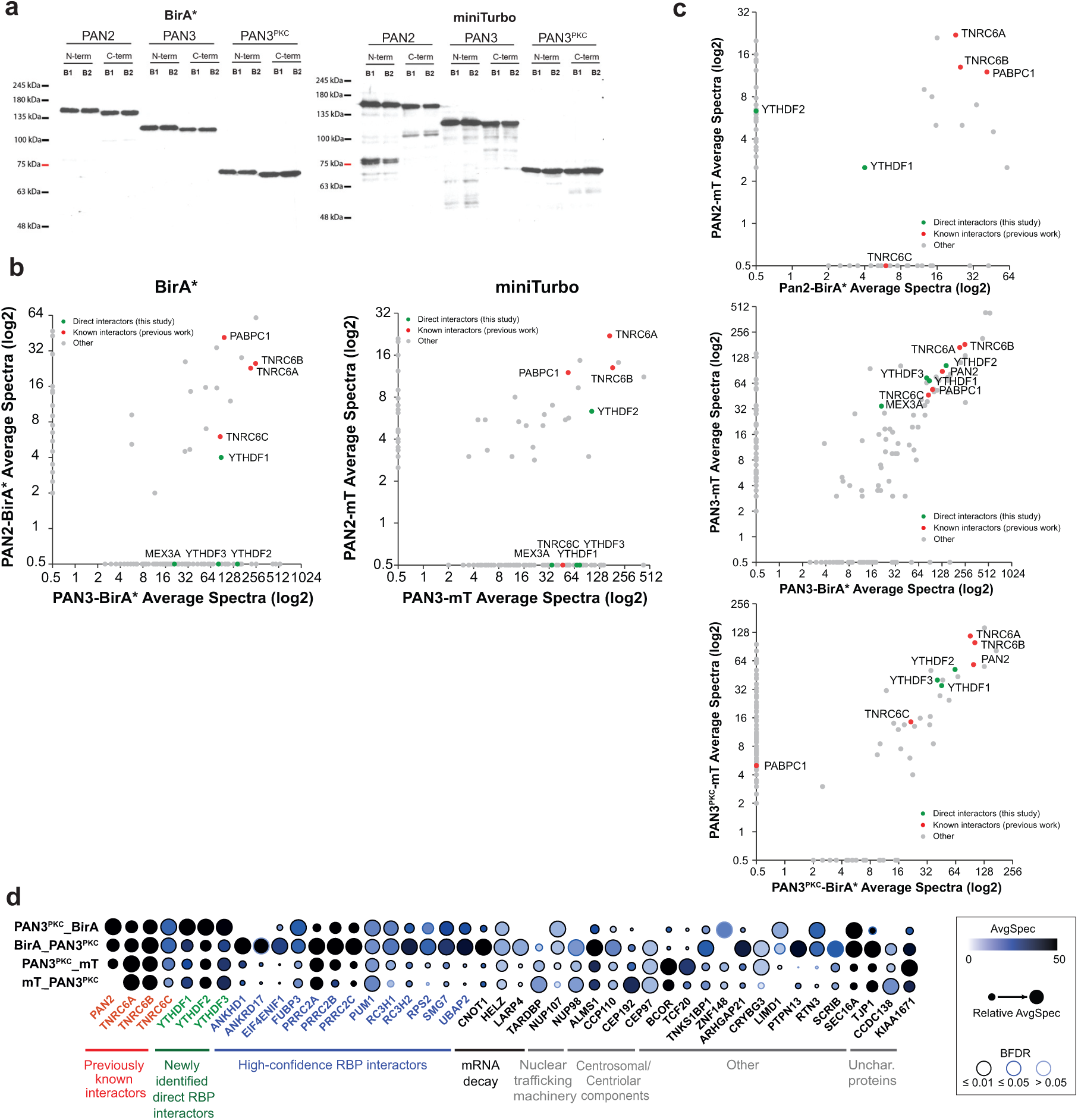
Identification of RNA-binding protein interactors of PAN2–PAN3 by proximity labelling. (a) **Western blot of tagged PAN2, PAN3 or PAN3^PKC^.** Each protein construct was cloned with N- or C-terminal BirA* or miniTurbo (mT) biotin ligase, in tandem with a FLAG tag, and transfected in duplicate into HEK293 cells. Similar expression levels were confirmed by detection of bands at the expected molecular weight using an anti-FLAG antibody. **(b) Proximity based proteomics of PAN2–PAN3.** A bait-bait scatter plot comparing Bio-ID of PAN2 (y-axis) and PAN3 (x-axis) tagged by BirA* (left) or miniTurbo (right), each averaged over the N- and C-terminal tags and filtered (PAN2-BirA*: n= 70; PAN2-mT: n=75; PAN3-BirA*: n=124; PAN3-mT: n=124); each dot represents a prey protein. High-confidence interactors of both PAN2 and PAN3 (BirA*: n=21; miniTurbo: n=27) lie within the graph body. All shown prey proteins have BFDR ≤ 0.01. Red: previously known interactors of PAN2–PAN3; green: direct interactors of PAN2–PAN3 characterized in this study. **(c) Interactors of PAN2–PAN3 identified through proximity labelling.** Bait-bait scatter plots comparing N- and C-terminally averaged miniTurbo- (y-axis) and BirA*-tagged (x-axis) PAN2 (top), PAN3 (middle), and PAN3^PKC^ (bottom); each dot represents a prey protein. For each tag, high-confidence prey proteins (BFDR ≤ 0.01) were obtained (PAN2-mT: n=75, PAN2-BirA: n=70; PAN3-mT: n=124; PAN3-BirA: n=124; PAN3^PKC^-mT: n=155; PAN3^PKC^-BirA: n=58). High-confidence interactors of each protein lie within the graph body (PAN2: n=13; PAN3: n=70; PAN3^PKC^: n=29). Red: previously known interactors of PAN2–PAN3; green: direct interactors of PAN2–PAN3 characterized in this study. **(d) Annotated dot plot of high-confidence interactors of PAN3^PKC^.** Data for PAN3^PKC^ tagged at the N- or C-terminus by BirA* (BirA-PAN3^PKC^, PAN3^PKC^-BirA) or miniTurbo (mT-PAN3^PKC^, PAN3^PKC^-mT) and used in proximity labelling assays (rows). Selected prey proteins were annotated by color for function as indicated. The dot plot is annotated by averaged spectral count (light blue to black), relative averaged spectral count between groups (dot size), and BFDR (outer circle color).

## References

1. Absmeier E, Chandrasekaran V, O’Reilly FJ, Stowell JAW, Rappsilber J, Passmore LA. 2023. Specific recognition and ubiquitination of translating ribosomes by mammalian CCR4-NOT. Nat Struct Mol Biol 30: 1314–1322.

2. Arvola RM, Chang C-T, Buytendorp JP, Levdansky Y, Valkov E, Freddolino PL, Goldstrohm AC. 2020. Unique repression domains of Pumilio utilize deadenylation and decapping factors to accelerate destruction of target mRNAs. Nucleic Acids Research 48: 1843–1871.

3. Ayed A, Mulder FA, Yi GS, Lu Y, Kay LE, Arrowsmith CH. 2001. Latent and active p53 are identical in conformation. Nat Struct Biol 8: 756–760.

4. Baltz AG, Munschauer M, Schwanhäusser B, Vasile A, Murakawa Y, Schueler M, Youngs N, Penfold-Brown D, Drew K, Milek M, et al. 2012. The mRNA-Bound Proteome and Its Global Occupancy Profile on Protein-Coding Transcripts. Molecular Cell 46: 674–690.

5. Basu S, Bahadur RP. 2016. A structural perspective of RNA recognition by intrinsically disordered proteins. Cell Mol Life Sci 73: 4075–4084.

6. Bhandari D, Raisch T, Weichenrieder O, Jonas S, Izaurralde E. 2014. Structural basis for the Nanos-mediated recruitment of the CCR4–NOT complex and translational repression. Genes Dev 28: 888–901.

7. Boeck R, Tarun S, Rieger M, Deardorff JA, Müller-Auer S, Sachs AB. 1996. The yeast Pan2 protein is required for poly(A)-binding protein-stimulated poly(A)-nuclease activity. J Biol Chem 271: 432–438.

8. Bönisch C, Temme C, Moritz B, Wahle E. 2007. Degradation of hsp70 and Other mRNAs in Drosophila via the 5′–3′ Pathway and Its Regulation by Heat Shock *. Journal of Biological Chemistry 282: 21818–21828.

9. Brandt T, Kaar JL, Fersht AR, Veprintsev DB. 2012. Stability of p53 Homologs. PLOS ONE 7: e47889.

10. Braun JE, Huntzinger E, Fauser M, Izaurralde E. 2011. GW182 proteins directly recruit cytoplasmic deadenylase complexes to miRNA targets. Mol Cell 44: 120–133.

11. Brown CE, Tarun SZ, Boeck R, Sachs AB. 1996. PAN3 encodes a subunit of the Pab1p-dependent poly(A) nuclease in Saccharomyces cerevisiae. Mol Cell Biol 16: 5744–5753.

12. Cano F, Bye H, Duncan LM, Buchet-Poyau K, Billaud M, Wills MR, Lehner PJ. 2012. The RNA-binding E3 ubiquitin ligase MEX-3C links ubiquitination with MHC-I mRNA degradation. EMBO J 31: 3596–3606.

13. Cano F, Rapiteanu R, Sebastiaan Winkler G, Lehner PJ. 2015. A non-proteolytic role for ubiquitin in deadenylation of MHC-I mRNA by the RNA-binding E3-ligase MEX-3C. Nat Commun 6: 8670.

14. Castello A, Fischer B, Eichelbaum K, Horos R, Beckmann BM, Strein C, Davey NE, Humphreys DT, Preiss T, Steinmetz LM, et al. 2012. Insights into RNA Biology from an Atlas of Mammalian mRNA-Binding Proteins. Cell 149: 1393–1406.

15. Chao H, Deng L, Xu F, Yu Z, Xu X, Huang J, Zeng T. 2019. MEX3C regulates lipid metabolism to promote bladder tumorigenesis through JNK pathway. Onco Targets Ther 12: 3285–3294.

16. Chen C-YA, Zheng D, Xia Z, Shyu A-B. 2009. Ago-TNRC6 triggers microRNA-mediated decay by promoting two deadenylation steps. Nat Struct Mol Biol 16: 1160–1166.

17. Cheng J, Maier KC, Avsec Ž, Rus P, Gagneur J. 2017. Cis-regulatory elements explain most of the mRNA stability variation across genes in yeast. RNA 23: 1648–1659.

18. Christie M, Boland A, Huntzinger E, Weichenrieder O, Izaurralde E. 2013. Structure of the PAN3 Pseudokinase Reveals the Basis for Interactions with the PAN2 Deadenylase and the GW182 Proteins. Molecular Cell 51: 360–373.

19. Coller J, Parker R. 2004. Eukaryotic mRNA decapping. Annu Rev Biochem 73: 861–890.

20. Draper BW, Mello CC, Bowerman B, Hardin J, Priess JR. 1996. MEX-3 is a KH domain protein that regulates blastomere identity in early C. elegans embryos. Cell 87: 205– 216.

21. Du H, Zhao Y, He J, Zhang Y, Xi H, Liu M, Ma J, Wu L. 2016. YTHDF2 destabilizes m6A-containing RNA through direct recruitment of the CCR4–NOT deadenylase complex. Nat Commun 7: 12626.

22. Eng JK, Jahan TA, Hoopmann MR. 2013. Comet: an open-source MS/MS sequence database search tool. Proteomics 13: 22–24.

23. Essig K, Kronbeck N, Guimaraes JC, Lohs C, Schlundt A, Hoffmann A, Behrens G, Brenner S, Kowalska J, Lopez-Rodriguez C, et al. 2018. Roquin targets mRNAs in a 3′-UTR-specific manner by different modes of regulation. Nat Commun 9: 3810.

24. Eulalio A, Tritschler F, Izaurralde E. 2009. The GW182 protein family in animal cells: New insights into domains required for miRNA-mediated gene silencing. RNA 15: 1433–1442.

25. Fabian MR, Frank F, Rouya C, Siddiqui N, Lai WS, Karetnikov A, Blackshear PJ, Nagar B, Sonenberg N. 2013. Structural basis for the recruitment of the human CCR4-NOT deadenylase complex by tristetraprolin. Nat Struct Mol Biol 20: 735–739.

26. Fox CA, Wickens M. 1990. Poly(A) removal during oocyte maturation: a default reaction selectively prevented by specific sequences in the 3’ UTR of certain maternal mRNAs. Genes Dev 4: 2287–2298.

27. Friedel CC, Dölken L, Ruzsics Z, Koszinowski UH, Zimmer R. 2009. Conserved principles of mammalian transcriptional regulation revealed by RNA half-life. Nucleic Acids Res 37: e115.

28. Garneau NL, Sokoloski KJ, Opyrchal M, Neff CP, Wilusz CJ, Wilusz J. 2008. The 3’ untranslated region of sindbis virus represses deadenylation of viral transcripts in mosquito and Mammalian cells. J Virol 82: 880–892.

29. Gaudet P, Livstone MS, Lewis SE, Thomas PD. 2011. Phylogenetic-based propagation of functional annotations within the Gene Ontology consortium. Brief Bioinform 12: 449–462.

30. Geertsma ER, Dutzler R. 2011. A Versatile and Efficient High-Throughput Cloning Tool for Structural Biology. Biochemistry 50: 3272–3278.

31. Geiger T, Wehner A, Schaab C, Cox J, Mann M. 2012. Comparative proteomic analysis of eleven common cell lines reveals ubiquitous but varying expression of most proteins. Mol Cell Proteomics 11: M111.014050.

32. Geisberg JV, Moqtaderi Z, Fan X, Ozsolak F, Struhl K. 2014. Global analysis of mRNA isoform half-lives reveals stabilizing and destabilizing elements in yeast. Cell 156: 812–824.

33. Goldstrohm AC, Hall TMT, McKenney KM. 2018. Post-transcriptional Regulatory Functions of Mammalian Pumilio Proteins. Trends Genet 34: 972–990.

34. Hao S, Baltimore D. 2009. The stability of mRNA influences the temporal order of the induction of genes encoding inflammatory molecules. Nat Immunol 10: 281–288.

35. Hill CH, Boreikaitė V, Kumar A, Casañal A, Kubík P, Degliesposti G, Maslen S, Mariani A, von Loeffelholz O, Girbig M, et al. 2019. Activation of the Endonuclease that Defines mRNA 3′ Ends Requires Incorporation into an 8-Subunit Core Cleavage and Polyadenylation Factor Complex. Mol Cell 73: 1217–1231.e11.

36. Hipps DS, Packman LC, Allen MD, Fuller C, Sakaguchi K, Appella E, Perham RN. 1994. The peripheral subunit-binding domain of the dihydrolipoyl acetyltransferase component of the pyruvate dehydrogenase complex of Bacillus stearothermophilus: preparation and characterization of its binding to the dihydrolipoyl dehydrogenase component. Biochem J 297 (**Pt 1**): 137–143.

37. Huang L, Malu S, McKenzie JA, Andrews MC, Talukder AH, Tieu T, Karpinets T, Haymaker C, Forget M-A, Williams LJ, et al. 2018. The RNA-binding Protein MEX3B Mediates Resistance to Cancer Immunotherapy by Downregulating HLA-A Expression. Clin Cancer Res 24: 3366–3376.

38. Hulsen T, de Vlieg J, Alkema W. 2008. BioVenn - a web application for the comparison and visualization of biological lists using area-proportional Venn diagrams. BMC Genomics 9: 488.

39. Huntzinger E, Kuzuoğlu-Öztürk D, Braun JE, Eulalio A, Wohlbold L, Izaurralde E. 2013. The interactions of GW182 proteins with PABP and deadenylases are required for both translational repression and degradation of miRNA targets. Nucleic Acids Res 41: 978–994.

40. Jasinski-Bergner S, Steven A, Seliger B. 2020. The Role of the RNA-Binding Protein Family MEX-3 in Tumorigenesis. Int J Mol Sci 21: 5209.

41. Jiao Y, Bishop CE, Lu B. 2012. Mex3c regulates insulin-like growth factor 1 (IGF1) expression and promotes postnatal growth. Mol Biol Cell 23: 1404–1413.

42. Jonas S, Izaurralde E. 2013. The role of disordered protein regions in the assembly of decapping complexes and RNP granules. Genes Dev 27: 2628–2641.

43. Jones DT, Cozzetto D. 2015. DISOPRED3: precise disordered region predictions with annotated protein-binding activity. Bioinformatics 31: 857–863.

44. Knight JDR, Choi H, Gupta GD, Pelletier L, Raught B, Nesvizhskii AI, Gingras A-C. 2017. ProHits-viz: a suite of web tools for visualizing interaction proteomics data. Nat Methods 14: 645–646.

45. Kozlov G, Ménade M, Rosenauer A, Nguyen L, Gehring K. 2010. Molecular determinants of PAM2 recognition by the MLLE domain of poly(A)-binding protein. J Mol Biol 397: 397–407.

46. Kuzuoglu-Öztürk D, Huntzinger E, Schmidt S, Izaurralde E. 2012. The Caenorhabditis elegans GW182 protein AIN-1 interacts with PAB-1 and subunits of the PAN2-PAN3 and CCR4-NOT deadenylase complexes. Nucleic Acids Res 40: 5651–5665.

47. Lee W, Tonelli M, Markley JL. 2015. NMRFAM-SPARKY: enhanced software for biomolecular NMR spectroscopy. Bioinformatics 31: 1325–1327.

48. Lin C, Miles WO. 2019. Beyond CLIP: advances and opportunities to measure RBP–RNA and RNA–RNA interactions. Nucleic Acids Research 47: 5490–5501.

49. Liu G, Zhang J, Larsen B, Stark C, Breitkreutz A, Lin Z-Y, Breitkreutz B-J, Ding Y, Colwill K, Pasculescu A, et al. 2010. ProHits: integrated software for mass spectrometry-based interaction proteomics. Nat Biotechnol 28: 1015–1017.

50. Liu J, Gao M, Xu S, Chen Y, Wu K, Liu H, Wang J, Yang X, Wang J, Liu W, et al. 2020. YTHDF2/3 Are Required for Somatic Reprogramming through Different RNA Deadenylation Pathways. Cell Rep 32: 108120.

51. Luo S, Tong L. 2014. Molecular basis for the recognition of methylated adenines in RNA by the eukaryotic YTH domain. Proc Natl Acad Sci U S A 111: 13834–13839.

52. Mangus DA, Evans MC, Agrin NS, Smith M, Gongidi P, Jacobson A. 2004. Positive and Negative Regulation of Poly(A) Nuclease. Mol Cell Biol 24: 5521–5533.

53. Mangus DA, Evans MC, Jacobson A. 2003. Poly(A)-binding proteins: multifunctional scaffolds for the post-transcriptional control of gene expression. Genome Biology 4: 223.

54. Mulder FA, Schipper D, Bott R, Boelens R. 1999. Altered flexibility in the substrate-binding site of related native and engineered high-alkaline Bacillus subtilisins. J Mol Biol 292: 111–123.

55. Notredame C, Higgins DG, Heringa J. 2000. T-Coffee: A novel method for fast and accurate multiple sequence alignment. J Mol Biol 302: 205–217.

56. Pagano JM, Farley BM, Essien KI, Ryder SP. 2009. RNA recognition by the embryonic cell fate determinant and germline totipotency factor MEX-3. Proceedings of the National Academy of Sciences 106: 20252–20257.

57. Parker R, Song H. 2004. The enzymes and control of eukaryotic mRNA turnover. Nat Struct Mol Biol 11: 121–127.

58. Passmore LA, Coller J. 2022. Roles of mRNA poly(A) tails in regulation of eukaryotic gene expression. Nat Rev Mol Cell Biol 23: 93–106.

59. Pekovic F, Lai WS, Corbo J, Hicks SN, Luke K, Blackshear PJ, Valkov E. 2025. Multivalent interactions with CCR4–NOT and PABPC1 determine mRNA repression efficiency by tristetraprolin. Nat Commun 16: 7528.

60. Plass M, Rasmussen SH, Krogh A. 2017. Highly accessible AU-rich regions in 3’ untranslated regions are hotspots for binding of regulatory factors. PLOS Computational Biology 13: e1005460.

61. Raisch T, Bhandari D, Sabath K, Helms S, Valkov E, Weichenrieder O, Izaurralde E. 2016. Distinct modes of recruitment of the CCR4-NOT complex by Drosophila and vertebrate Nanos. EMBO J 35: 974–990.

62. Robert X, Gouet P. 2014. Deciphering key features in protein structures with the new ENDscript server. Nucleic Acids Res 42: W320–324.

63. Sandler H, Kreth J, Timmers HTM, Stoecklin G. 2011. Not1 mediates recruitment of the deadenylase Caf1 to mRNAs targeted for degradation by tristetraprolin. Nucleic Acids Res 39: 4373–4386.

64. Schaefer JS, Klein JR. 2016. Roquin--a multifunctional regulator of immune homeostasis. Genes Immun 17: 79–84.

65. Schäfer IB, Yamashita M, Schuller JM, Schüssler S, Reichelt P, Strauss M, Conti E. 2019. Molecular Basis for poly(A) RNP Architecture and Recognition by the Pan2-Pan3 Deadenylase. Cell 177: 1619–1631.e21.

66. Schueler M, Munschauer M, Gregersen LH, Finzel A, Loewer A, Chen W, Landthaler M, Dieterich C. 2014. Differential protein occupancy profiling of the mRNA transcriptome. Genome Biology 15: R15.

67. Sgromo A, Raisch T, Bawankar P, Bhandari D, Chen Y, Kuzuoğlu-Öztürk D, Weichenrieder O, Izaurralde E. 2017. A CAF40-binding motif facilitates recruitment of the CCR4-NOT complex to mRNAs targeted by Drosophila Roquin. Nat Commun 8: 14307.

68. Silverman IM, Li F, Alexander A, Goff L, Trapnell C, Rinn JL, Gregory BD. 2014. RNase-mediated protein footprint sequencing reveals protein-binding sites throughout the human transcriptome. Genome Biology 15: R3.

69. Stowell JAW, Webster MW, Kögel A, Wolf J, Shelley KL, Passmore LA. 2016. Reconstitution of Targeted Deadenylation by the Ccr4-Not Complex and the YTH Domain Protein Mmi1. Cell Rep 17: 1978–1989.

70. Stowell JAW, Yu CWH, Chen ZA, Lee G, Morgan T, Sinn L, Agnello S, O’Reilly FJ, Rappsilber J, Freund SMV, et al. 2024. Phosphorylation-dependent tuning of mRNA deadenylation rates. bioRxiv 2024.10.18.618793.

71. Teo G, Liu G, Zhang J, Nesvizhskii AI, Gingras A-C, Choi H. 2014. SAINTexpress: improvements and additional features in Significance Analysis of INTeractome software. J Proteomics 100: 37–43.

72. Theler D, Dominguez C, Blatter M, Boudet J, Allain FH-T. 2014. Solution structure of the YTH domain in complex with N6-methyladenosine RNA: a reader of methylated RNA. Nucleic Acids Res 42: 13911–13919.

73. Tompa P, Davey NE, Gibson TJ, Babu MM. 2014. A Million Peptide Motifs for the Molecular Biologist. Molecular Cell 55: 161–169.

74. Tucker M, Staples RR, Valencia-Sanchez MA, Muhlrad D, Parker R. 2002. Ccr4p is the catalytic subunit of a Ccr4p/Pop2p/Notp mRNA deadenylase complex in Saccharomyces cerevisiae. EMBO J 21: 1427–1436.

75. Tucker M, Valencia-Sanchez MA, Staples RR, Chen J, Denis CL, Parker R. 2001. The Transcription Factor Associated Ccr4 and Caf1 Proteins Are Components of the Major Cytoplasmic mRNA Deadenylase in Saccharomyces cerevisiae. Cell 104: 377– 386.

76. Uchida N, Hoshino S, Katada T. 2004. Identification of a Human Cytoplasmic Poly(A) Nuclease Complex Stimulated by Poly(A)-binding Protein*. Journal of Biological Chemistry 279: 1383–1391.

77. Van Nostrand EL, Freese P, Pratt GA, Wang X, Wei X, Xiao R, Blue SM, Chen J-Y, Cody NAL, Dominguez D, et al. 2020a. A large-scale binding and functional map of human RNA-binding proteins. Nature 583: 711–719.

78. Van Nostrand EL, Pratt GA, Yee BA, Wheeler EC, Blue SM, Mueller J, Park SS, Garcia KE, Gelboin-Burkhart C, Nguyen TB, et al. 2020b. Principles of RNA processing from analysis of enhanced CLIP maps for 150 RNA binding proteins. Genome Biology 21: 90.

79. Varadi M, Zsolyomi F, Guharoy M, Tompa P. 2015. Functional Advantages of Conserved Intrinsic Disorder in RNA-Binding Proteins. PLOS ONE 10: e0139731.

80. Wahle E, Winkler GS. 2013. RNA decay machines: deadenylation by the Ccr4-not and Pan2-Pan3 complexes. Biochim Biophys Acta 1829: 561–570.

81. Wang C, Uversky VN, Kurgan L. 2016. Disordered nucleiome: Abundance of intrinsic disorder in the DNA- and RNA-binding proteins in 1121 species from Eukaryota, Bacteria and Archaea. PROTEOMICS 16: 1486–1498.

82. Wang M, Herrmann CJ, Simonovic M, Szklarczyk D, von Mering C. 2015. Version 4.0 of PaxDb: Protein abundance data, integrated across model organisms, tissues, and cell-lines. Proteomics 15: 3163–3168.

83. Wang X, Lu Z, Gomez A, Hon GC, Yue Y, Han D, Fu Y, Parisien M, Dai Q, Jia G, et al. 2014. N6-methyladenosine-dependent regulation of messenger RNA stability. Nature 505: 117–120.

84. Wang Y, Liu CL, Storey JD, Tibshirani RJ, Herschlag D, Brown PO. 2002. Precision and functional specificity in mRNA decay. Proc Natl Acad Sci U S A 99: 5860–5865.

85. Webster MW, Stowell JA, Passmore LA. 2019. RNA-binding proteins distinguish between similar sequence motifs to promote targeted deadenylation by Ccr4-Not. Elife 8: e40670.

86. Webster MW, Stowell JAW, Tang TTL, Passmore LA. 2017. Analysis of mRNA deadenylation by multi-protein complexes. Methods 126: 95–104.

87. Weissmann F, Petzold G, VanderLinden R, Huis in ’t Veld PJ, Brown NG, Lampert F, Westermann S, Stark H, Schulman BA, Peters J-M. 2016. biGBac enables rapid gene assembly for the expression of large multisubunit protein complexes. Proceedings of the National Academy of Sciences 113: E2564–E2569.

88. Xu C, Wang X, Liu K, Roundtree IA, Tempel W, Li Y, Lu Z, He C, Min J. 2014. Structural basis for selective binding of m6A RNA by the YTHDC1 YTH domain. Nat Chem Biol 10: 927–929.

89. Yamashita A, Chang T-C, Yamashita Y, Zhu W, Zhong Z, Chen C-YA, Shyu A-B. 2005. Concerted action of poly(A) nucleases and decapping enzyme in mammalian mRNA turnover. Nat Struct Mol Biol 12: 1054–1063.

90. Yang L, Wang C, Li F, Zhang J, Nayab A, Wu J, Shi Y, Gong Q. 2017. The human RNA-binding protein and E3 ligase MEX-3C binds the MEX-3–recognition element (MRE) motif with high affinity. J Biol Chem 292: 16221–16234.

91. Yi H, Park J, Ha M, Lim J, Chang H, Kim VN. 2018. PABP Cooperates with the CCR4-NOT Complex to Promote mRNA Deadenylation and Block Precocious Decay. Molecular Cell 70: 1081–1088.e5.

92. Youn J-Y, Dunham WH, Hong SJ, Knight JDR, Bashkurov M, Chen GI, Bagci H, Rathod B, MacLeod G, Eng SWM, et al. 2018. High-Density Proximity Mapping Reveals the Subcellular Organization of mRNA-Associated Granules and Bodies. Mol Cell 69: 517–532.e11.

93. Zhao B, Katuwawala A, Oldfield CJ, Hu G, Wu Z, Uversky VN, Kurgan L. 2021. Intrinsic Disorder in Human RNA-Binding Proteins. Journal of Molecular Biology 433: 167229.

