## Supplementary Information 1 for "RNA-binding proteins provide specificity to the PAN2–PAN3 mRNA deadenylation complex"

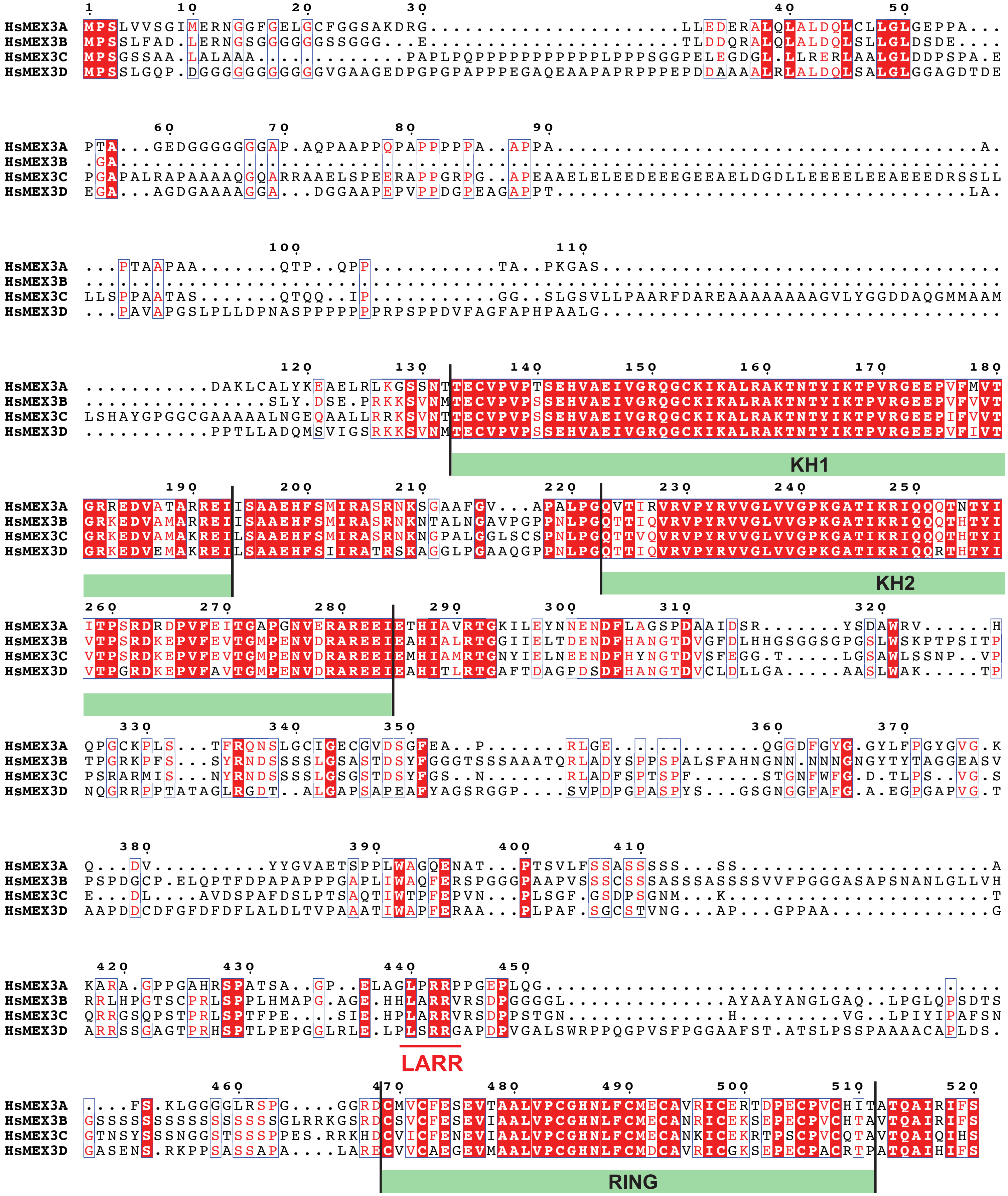


**Supplementary Information 1: Sequence alignment of human MEX3 paralogs (MEX3A-D).** Residues identical in all four paralogs are in white with a red background, similar residues are in red and framed in blue. Domain boundaries are indicated by black lines and green boxes, and the LARR motif is underlined.
