## Supplementary Information 2 for "RNA-binding proteins provide specificity to the PAN2–PAN3 mRNA deadenylation complex"

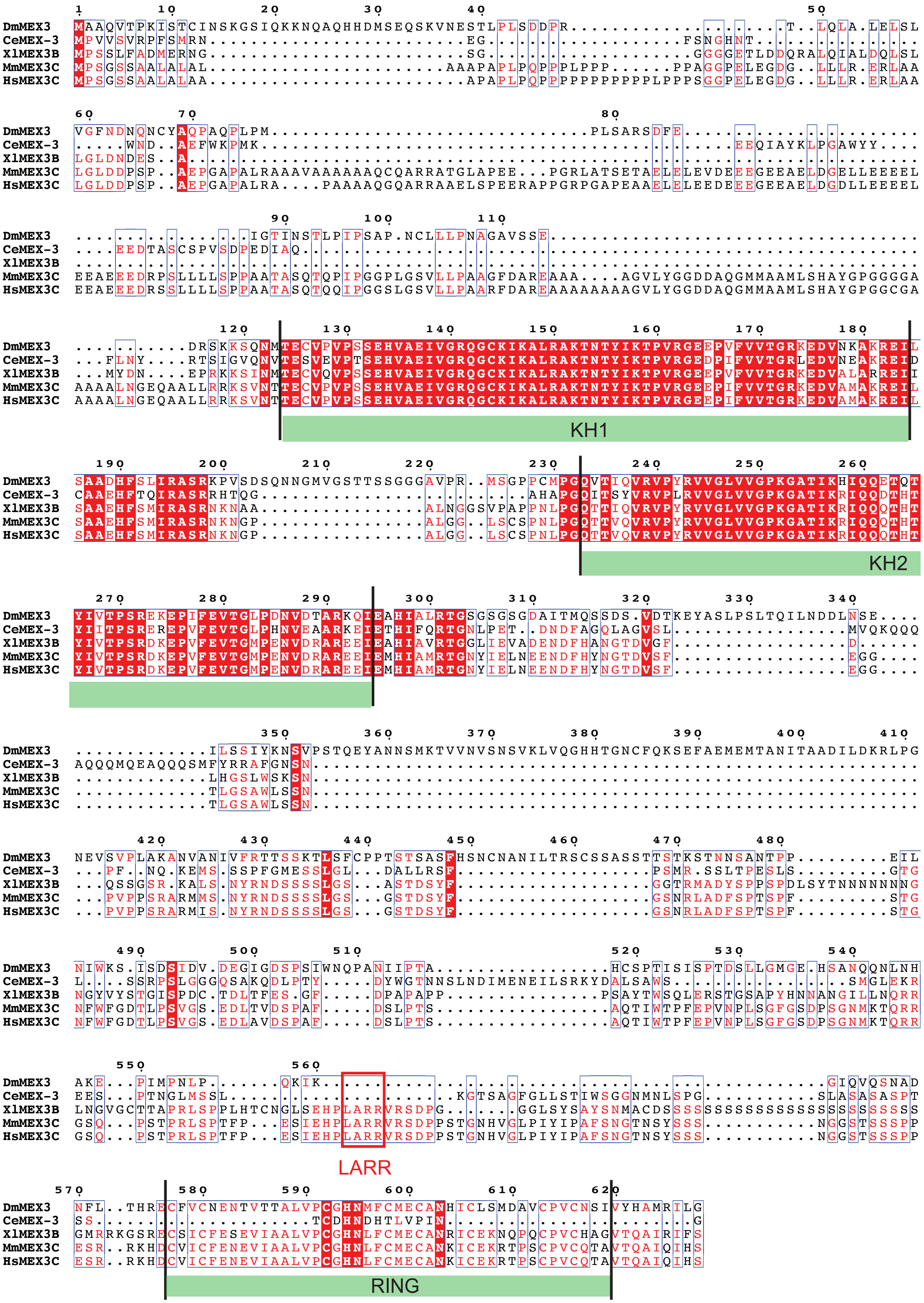


**Supplementary Information 2: Sequence alignment of MEX3 among selected species.** Dm: *Drosophila melanogaster*; Ce: *Caenorhabditis elegans*; Xl: *Xenopus laevis*; Mm: *Mus musculus*; Hs: *Homo sapiens*. The LARR motif is boxed in red, and is conserved in *Xenopus laevis*, *Mus musculus*, and *Homo sapiens*. Fully conserved residues across all five species are in white with a red background; mostly (>60%) conserved residues (found in three species) are in red with a blue box. The RNA-binding tandem KH domains and RING domain (green bars) are well-conserved, except in *C. elegans*.
