## Supplementary Information 3 for "RNA-binding proteins provide specificity to the PAN2–PAN3 mRNA deadenylation complex"

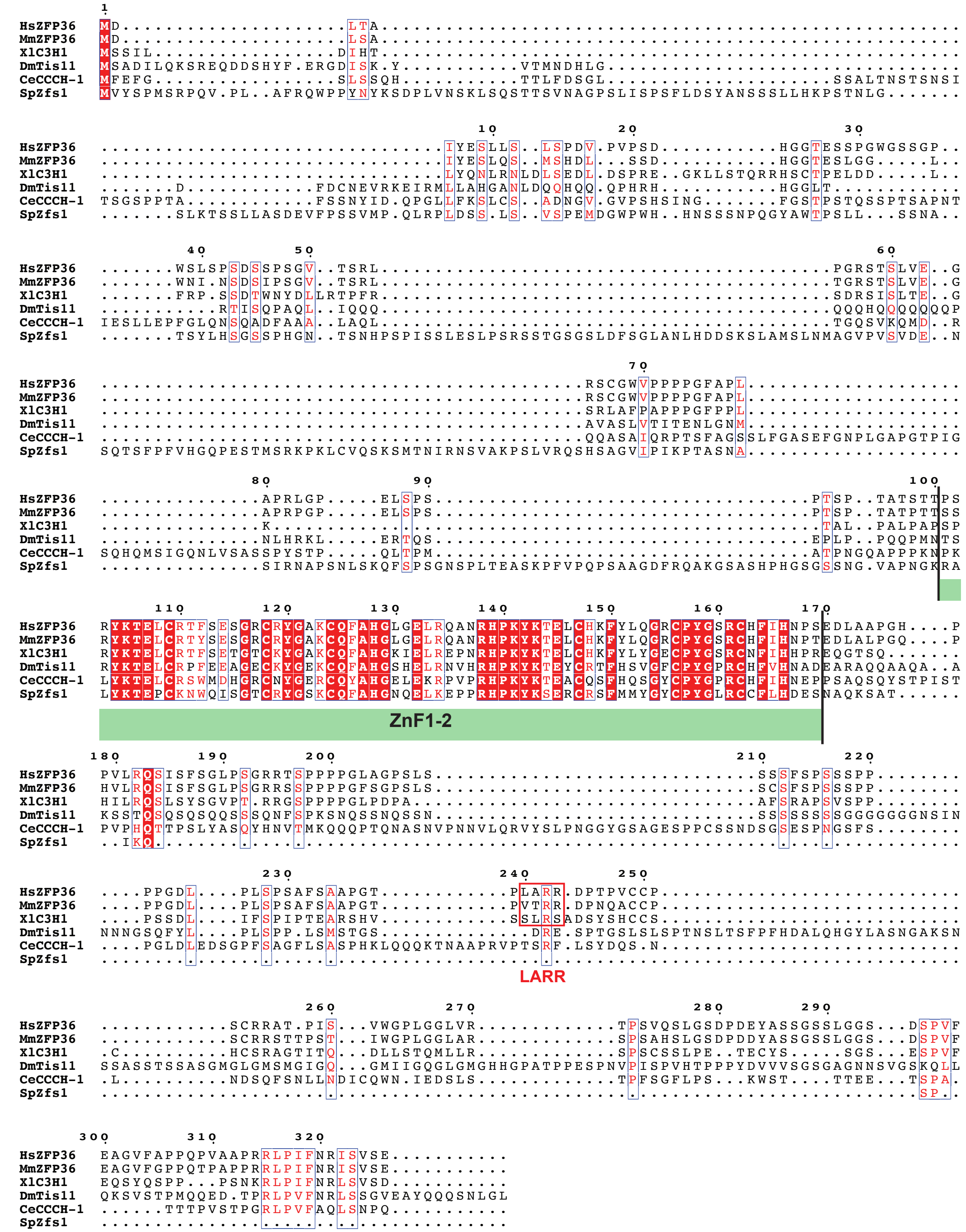


**Supplementary Information 3: Conservation of the IARK motif of YTHDF3 in vertebrates.** Sequence alignment of YTHDF3 from *Homo sapiens* (Hs), *Mus musculus* (Mm), *Xenopus laevis* (Xl), and YTHDF from *Drosophila melanogaster* (Dm), and *Saccharomyces cerevisiae* (ScMRB1). The YTH domain (green bar) is conserved among YTHDF3 homologs. The IARK motif (boxed in red) is conserved in vertebrates, including *Homo sapiens*, *Mus musculus* and *Xenopus laevis*.
