## Supplementary Information 4 for "RNA-binding proteins provide specificity to the PAN2–PAN3 mRNA deadenylation complex"

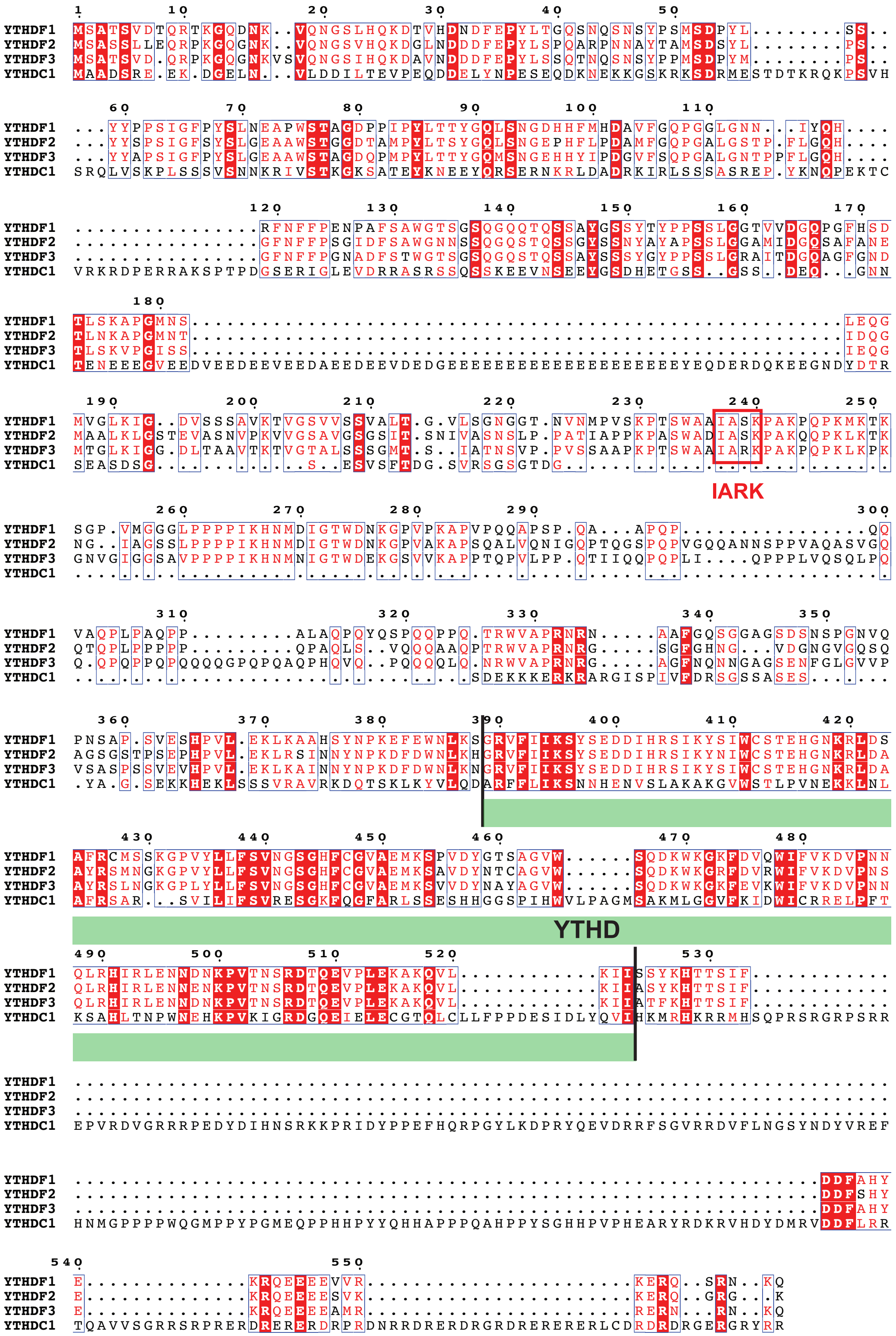


**Supplementary Information 4: Sequence alignment of human YTHDF1, YTHDF2, YTHDF3, and YTHDC1.** The YTH domain (residues 389-523; green bar) is highly conserved among YTHDF and YTHDC proteins. The IARK motif (boxed in red) is absent in YTHDC1, consistent with the lack of a known role in cytoplasmic post-transcriptional gene regulation. In YTHDF1 and YTHDF2, the IARK motif is replaced by an IASK motif. Conserved residues are in white with a red background; mostly conserved residues (>75%) are in red with a white box.
