## Supplementary Information 5 for "RNA-binding proteins provide specificity to the PAN2–PAN3 mRNA deadenylation complex"

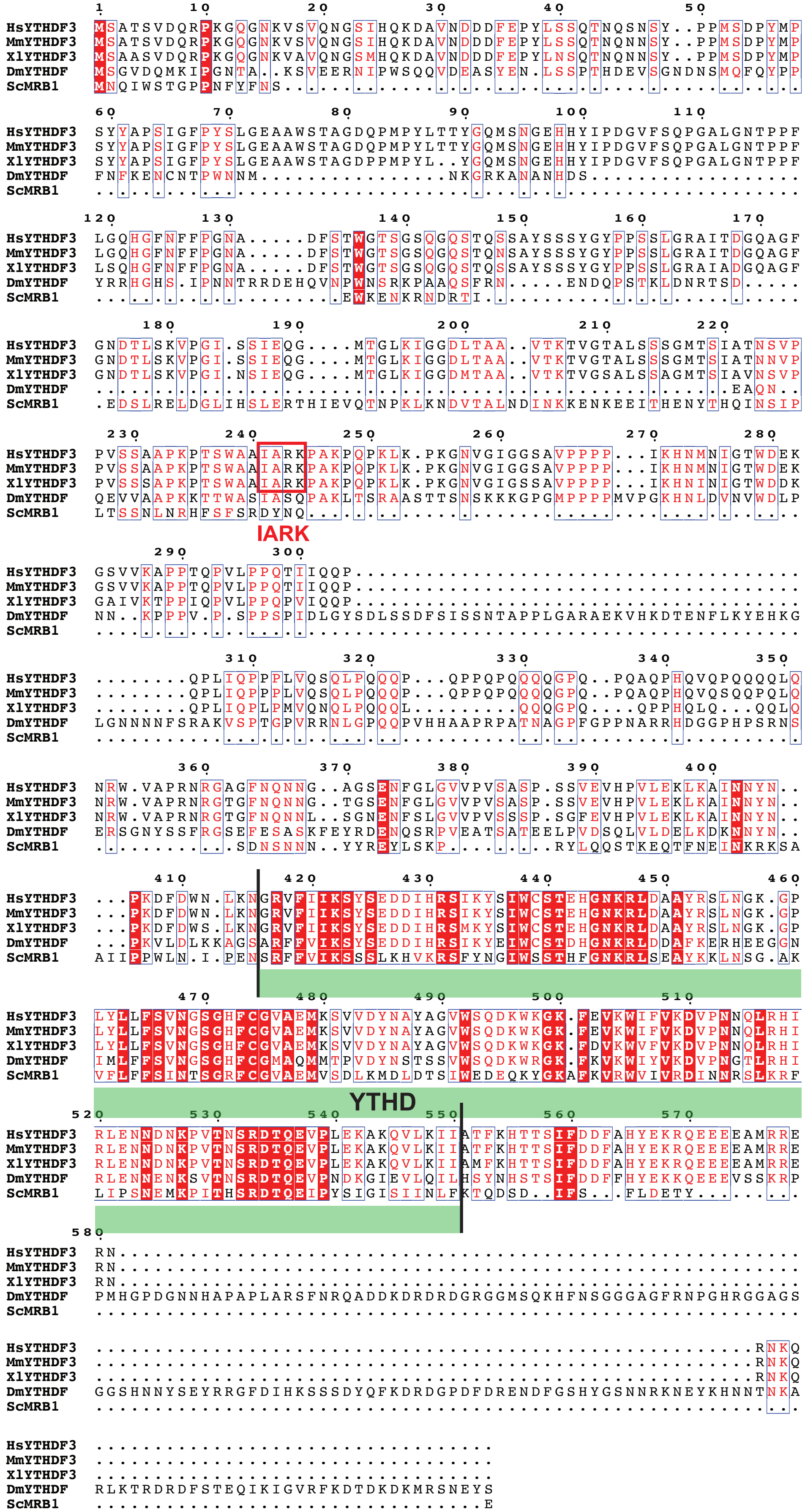


**Supplementary Information 5: Sequence conservation of ZFP36 LARR-like motif in different species.** Sequence alignment of ZFP36/TTP in *Homo sapiens* (Hs), *Mus musculus* (Mm), *Xenopus laevis* (Xl), *Drosophila melanogaster* (Dm), *Caenorhabditis elegans* (Ce), and *Saccharomyces cerevisiae* (Sc). Conserved residues are in white with a red background; mostly conserved residues (>66%) are in red with a white box. The RNA-binding zinc finger domains (ZnF1-2, green bar) are well conserved among ZFP36 homologs. The LARR motif (boxed in red) is conserved only in mammals; in mouse, the LARR motif is replaced by VTRR.
